# Dynamin is primed at endocytic sites for ultrafast endocytosis

**DOI:** 10.1101/2021.02.15.431332

**Authors:** Yuuta Imoto, Sumana Raychaudhuri, Pascal Fenske, Eduardo Sandoval, Kie Itoh, Eva-Maria Blumrich, Lauren Mamer, Fereshteh Zarebidaki, Berit Söhl-Kielczynski, Thorsten Trimbuch, Shraddha Nayak, Janet H. Iwasa, Erik M. Jorgensen, Michael A. Cousin, Christian Rosenmund, Shigeki Watanabe

## Abstract

Dynamin mediates fission of vesicles from the plasma membrane during endocytosis. Typically, dynamin is recruited from the cytosol to endocytic sites, requiring seconds to tens of seconds. However, ultrafast endocytosis in neurons internalizes vesicles as quickly as 50 ms during synaptic vesicle recycling. Here we demonstrate that Dynamin 1 is pre-recruited to endocytic sites for ultrafast endocytosis. Specifically, Dynamin 1xA, a splice variant of Dynamin 1, interacts with Syndapin 1 to form molecular condensates on the plasma membrane when the proline-rich domain of this variant is dephosphorylated. When this domain is mutated to include phosphomimetic residues or Syndapin 1’s dynamin-interacting domain is mutated, Dynamin 1xA becomes diffuse, and consequently, ultrafast endocytosis slows down by ∼100-fold. Mechanistically, Syndapin 1 acts as an adaptor by binding the plasma membrane and stores Dynamin 1xA at endocytic sites. This cache bypasses the recruitment step and accelerates endocytosis at synapses.

## Introduction

Dynamin GTPase (Shpetner and Vallee, 1989) catalyzes membrane fission during endocytosis (Praefcke and McMahon, 2004). Dynamin forms a contractile helix around the neck of endocytic pits and cleaves the neck upon GTP hydrolysis (Antonny et al., 2016; Hinshaw and Schmid, 1995; Praefcke and McMahon, 2004; Takei et al., 1995). *In vitro* experiments suggest that constriction of the membrane is rapid, and is completed within hundreds of milliseconds (Morlot et al., 2012). These experiments were conducted at saturated dynamin concentrations and unlimited GTP. However, live cell imaging suggests that the amount of dynamin around endocytic pits is initially low and increases only as pits mature during clathrin-mediated endocytosis. This increase is due to the gradual recruitment of dynamin from the cytosol, which requires ∼10–30 s (Cocucci et al., 2014; Taylor et al., 2011a, 2012), and is dependent on interactions with other endocytic proteins such as Endophilin A (Meinecke et al., 2013). Since dynamin interacting proteins are also being recruited from the cytosol (Taylor et al., 2011a, 2012), clathrin-mediated endocytosis is inherently slow, requiring seconds to minutes.

Despite these slow kinetics, dynamin is thought to play a key role in synaptic vesicle recycling. In neuromuscular junctions of *Drosophila melanogaster*, endocytic pits are arrested on the plasma membrane in animals carrying a temperature-sensitive allele of dynamin (*Shi^ts^*) following a heat shock, resulting in rapid depletion of synaptic vesicles (Koenig and Ikeda, 1989; Kosaka and Ikeda, 1983). Likewise, ultrastructural studies in *Caenorhabditis elegans* (*C. elegans*) and *Mus musculus* (mouse, hereafter) lacking functional dynamin show stalled endocytic pits at presynapses (Ferguson et al., 2007; Kittelmann et al., 2013; Raimondi et al., 2011). However, recent studies using time-resolved “flash-and-freeze” electron microscopy suggest that endocytosis at synapses is much faster (∼30 ms – 1 s) than typical clathrin-mediated endocytosis (Watanabe et al., 2013a, 2013b). During ultrafast endocytosis, synaptic vesicle exocytosis triggers the formation of endocytic pits next to the active zone where synaptic vesicles fuse, and these pits are internalized between 50 ms to 300 ms at mouse synapses. Following the delivery of endocytic vesicles to endosomes, clathrin-coated vesicles bud off from endosomes to regenerate synaptic vesicles (Watanabe et al., 2014). Ultrafast endocytosis depends on dynamin in *C. elegans*; endocytic pits are stuck on the plasma membrane at the restrictive temperature in temperature-sensitive mutants of dynamin (Clark et al., 1997) (Watanabe et al., 2013a). Similarly, ultrafast endocytosis is stalled in mouse hippocampal synapses after an application of the dynamin inhibitor Dynasore (Watanabe et al., 2013b), suggesting dynamin likely mediates ultrafast endocytosis. However, given the slow recruitment of dynamin and other endocytic proteins from the cytosol, it is not clear how dynamin can pinch off endocytic intermediates on this time scale.

In mammals, there are three isoforms of dynamin; Dynamin 1, 2, 3, encoded by DNM1, 2, 3, genes in mouse, respectively (Cook et al., 1996). All dynamin isoforms contain an N-terminal GTPase domain, a Pleckstrin homology (PH) domain, and a C-terminal proline-rich domain (PRD) (Cao et al., 1998). Dynamin 2 (or Dyn2) is ubiquitously expressed and essential for the development of a nervous system (Ferguson et al., 2009), while Dynamin 1 and 3 (or Dyn1,3) are highly expressed in brain and primarily involved in synaptic vesicle recycling (Ferguson et al., 2007; Raimondi et al., 2011).

Dynamin function is regulated by phosphorylation of the PRD. The PRD is phosphorylated by multiple kinases including GSK3ß (S774) and CDK5 (S778) and dephosphorylated by Calcineurin. Under resting conditions, approximately 75% of Dyn1 present at synaptic terminals is dephosphorylated (Graham et al., 2007; Liu et al., 1994), and this fraction is increased or decreased based on the activity level of neurons (Clayton et al., 2009). The dephosphorylated PRD of Dyn1 binds the Src-Homology 3 (SH3) domain of Bin, Amphyphysin, Rvs (BAR) proteins, Syndapin 1, Amphiphysin 1, and Endophilin A (Ferguson and De Camilli, 2012). The interaction between Dyn1 and Syndapin 1 or Endophlin A is reduced by phosphorylation of serines S774 and S778 within the PRD of dynamin (Anggono et al., 2006) (Xue et al., 2011). The endocytic defect in *Dyn1,3 DKO* can be rescued by the expression of wild-type Dyn1 or phospho-deficient Dyn1 (S774/778A) (Armbruster et al., 2013). However, endocytosis is slowed when a phosphomimetic form (S774/778D) is expressed (Armbruster et al., 2013), suggesting that the acceleration of endocytosis at synapses likely depends on these binding partners. However, even when Dyn1 is dephosphorylated, the time constant of endocytosis is ∼10 s (Armbruster et al., 2013), and thus, how dynamin mediates ultrafast endocytosis remains unclear.

Here we used a combination of flash-and-freeze electron microscopy and fluorescence microscopy in mouse hippocampal neurons to examine the roles of dynamin in ultrafast endocytosis. We found that deeply invaginated endocytic pits are stalled at the plasma membrane during ultrafast endocytosis in *Dyn1,3* double knockout (DKO) neurons. This defect can be rescued by the expression of a splice variant of Dyn1, Dyn1xA, but not another variant xB. Importantly, Dyn1xA is localized at putative endocytic sites at synapses due to multivalent interactions with Syndapin 1 and Endophilin A. When the expression of Syndapin 1 is knocked down or its membrane interacting domain is mutated (I122D/M123E), Dyn1xA becomes diffuse, and ultrafast endocytosis slows down 100-fold, suggesting that Syndapin 1 acts as an adaptor between the plasma membrane and dynamin. Similarly, the phosphomimetic form of Dyn1 (S774/778D) is not recruited to the plasma membrane, and consequently, endocytic pits persist on the plasma membrane for over 10 s. More specifically, the constriction of the neck of pits is slowed down under these conditions due to the slow recruitment of dynamin. Together, these results suggest that dynamin is pre-assembled at endocytic sites, bypassing the need for recruitment from the cytosol, to accelerate endocytosis at synapses.

## Results

### Dynamin 1 is required for ultrafast endocytosis

Ultrafast endocytosis is blocked by the dynamin inhibitor Dynasore (Watanabe et al., 2013b), indicating a possible involvement of dynamin. However, it is not clear which isoform of dynamin is necessary since Dynasore is not isoform-specific. In addition, Dynasore has off-target effects (Park et al., 2013; Yamada et al., 2009), making it uncertain whether dynamin is actually involved. To address these issues, we first performed flash-and-freeze experiments in neurons lacking the two brain-enriched isoforms of dynamin, Dyn1 and 3 (Figure 1). *DNM1^+/-^,3 ^-/-^* mice were bred to obtain *DNM1*^+/+^,*3*^-/-^ (*Dyn3* KO) and *DNM1*^-/-^,*3*^-/-^ (*Dyn1,3* DKO). *DNM1^+/-^,3 ^-/-^* mice were not used in this study. *Dyn3* KO served as a control since *Dyn3* KO itself does not show apparent functional or structural phenotypes (Ferguson et al., 2007; Raimondi et al., 2011), although it exacerbates the *Dyn1* KO phenotypes (Raimondi et al., 2011). Because *Dyn1,3* DKO (*DNM1*^-/-^,*3*^-/-^) mice die perinatally (Raimondi et al., 2011), we cultured hippocampal neurons from animals of embryonic days 18 (E18) or postnatal day 0 (P0) immediately after birth. These neurons were infected with lentivirus expressing a variant of channelrhodopsin, ChetaTC (Berndt et al., 2011; Gunaydin et al., 2010), and the experiments performed around 13-16 days *in vitro* (DIV13-16). To block an accumulation of endocytic intermediates in *Dyn1*,*3* DKO neurons due to spontaneous network activity, neurons were incubated with sodium channel blocker tetrodotoxin (TTX, 1 µM) overnight to silence neural activity – this treatment almost completely reverts the accumulation of endocytic intermediates (Raimondi et al., 2011; Wu et al., 2014). On the next day, TTX was washed off thoroughly, and a fluid phase marker, cationized ferritin, was applied in the external physiological saline (2 mg/ml, 5 min, 1 mM Ca^2+^). A 10-ms pulse of light was applied, and the subsequent membrane dynamics were captured by freezing the neurons at 1 s or 10 s after a single stimulus (37 °C, 4 mM Ca^2+^). These time points are chosen because the majority of ultrafast endocytosis complete by 1s – under normal conditions ferritin particles are in endocytic vesicles or endosomes by this time point and subsequently in synaptic vesicles by ∼10 s (Watanabe et al., 2013b, 2014).

**Figure 1.**
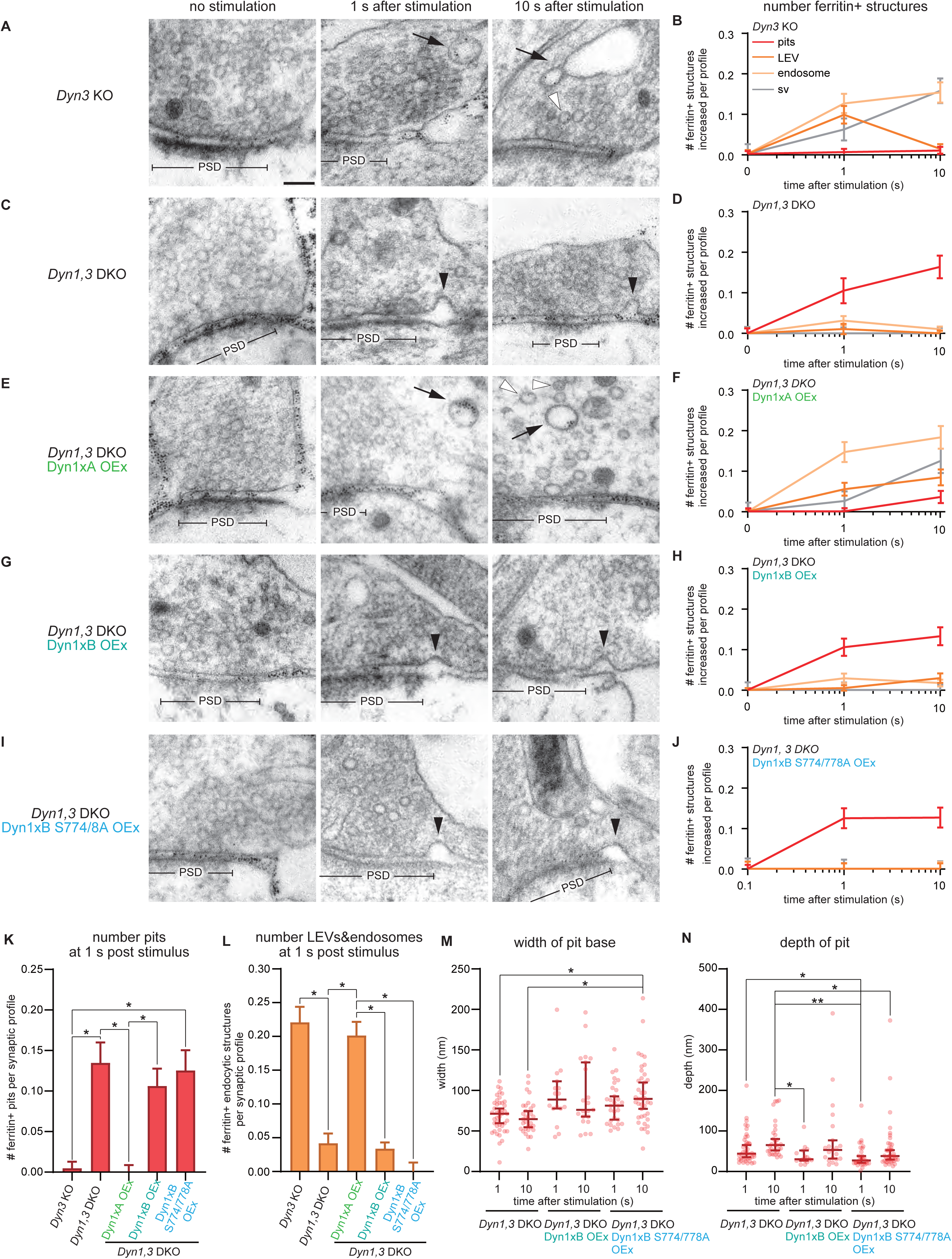
Dyn1xA splice variant, but not xB variant, mediates ultrafast endocytosis. (A, C, E, G and I) Example micrographs showing endocytic pits and ferritin-containing endocytic structures at the indicated time points in *Dyn3* KO (A), *Dyn1, 3* DKO (C), *Dyn1, 3* DKO, Dyn1xA overexpression (OEx) (E), *Dyn1, 3* DKO, Dyn1xB OEx (G), *Dyn1, 3* DKO, Dyn1xB S774/778A OEx (I). Black arrowheads, endocytic pits; black arrows, large endocytic vesicles (LEVs) or endosomes; white arrowheads, synaptic vesicles. Scale bar: 100 nm. PSD, post-synaptic density. (B, D, F, H, and J) Plots showing the increase in the number of each endocytic structure per synaptic profile after a single stimulus in *Dyn3* KO (B), *Dyn1, 3* DKO (D), *Dyn1, 3* DKO, Dyn1xA OEx (F), *Dyn1, 3* DKO and Dyn1xB OEx (H) and *Dyn1, 3* DKO, Dyn1xB S774/778A OEx (J). The mean and SEM are shown in each graph. (K) Number of endocytic pits at 1s after stimulation. The numbers are re-plotted as a bar graph from the 1 s time point in (B, D, F, H, and J) for easier comparison between groups. Ordinary one-way ANOVA with full pairwise comparisons by Holm-Šídák’s multiple comparisons test. The mean and SEM are shown. (L) Number of LEVs and endosomes at 1s after stimulation. The numbers of LEVs and endosomes are summed from the data presented in (B, D, F, H, and J), averaged, and re-plotted for easier comparison between groups. Ordinary one-way ANOVA with full pairwise comparisons by Holm-Šídák’s multiple comparisons test. The mean and SEM are shown. (M and N) Plots showing the width (M) and depth (N) of endocytic pits at the 1s time point. Mann-Whitney test was used compared each value to *Dyn1, 3* DKO at 1 s. Kruskal–Wallis tests with full comparisons by post hoc Dunn’s multiple comparisons tests. n = *Dyn1, 3* DKO, 42 pits for 1 s and 31 pits for 10 s; *Dyn1, 3* DKO, Dyn1xB OEx, 16 pits for 1 s and 21 pits for 10 s; and *Dyn1, 3* DKO, Dyn1xB S774/778A OEx, 29 pits for 1 s and 39 pits for 10 s. All data are from two independent experiments from N = 2 cultures prepared and frozen on different days. n = *Dyn* 3KO, 649; *Dyn1, 3* DKO, 622; *Dyn1, 3* DKO, Dyn1xA OEx, 642; *Dyn1, 3* DKO, Dyn1xB OEx, 666; and *Dyn1, 3* DKO, Dyn1xB S774/778A OEx, 614 synaptic profiles in (B, D, F, H, J, K, and L). *p < 0.05, **p < 0.0001. See Quantification and Statistical Analysis for the n values and detailed numbers for each time point. Knock out neurons are from the littermates in all cases.

Like in wild-type neurons (Watanabe et al., 2013b, 2014), ferritin particles were first observed in large endocytic vesicles and synaptic endosomes at 1 s after stimulation and then in synaptic vesicles at 10 s in *Dyn3* KO neurons (Figure 1A, 1B), suggesting that ultrafast endocytosis is intact and Dyn3 is likely dispensable for this process. However, ferritin-positive endosomes seemed to persist in these neurons, suggesting that Dyn3 may play a role in the resolution of synaptic endosomes (Figure 1A, 1B). By contrast, in *Dyn1,3* DKO neurons, endocytic pits were stuck at the plasma membrane (Figure 1C,1D, S1A) immediately next to the active zone, at the putative sites of ultrafast endocytosis (Fig. S1A, S1D, S1E), and no ferritin particles were observed in endocytic vesicles and endosomes at either time point (Figure 1C, 1D, 1K, 1L), suggesting that endocytic structures cannot be internalized in the DKO neurons. No electron density indicative of clathrin-coats was observed around these pits (Figure 1C, S1B, S1C). Occasionally, clathrin-coated pits were observed distant from the active zone (Figure S1B, S1D, translucent arrowhead), but the number did not increase following stimulation (Figure S1C) (Imig et al., 2020; Watanabe et al., 2013b, 2014). Endocytic pits in the DKO neurons were deeply invaginated (Figure 1N; pit depth, 44.2 ± 6.0 nm, n=42) and became further tubulated by 10 s (Figure 1N; pit depth, 65.0 ± 7.7 nm, n=31, p = 0.240). The width of the pit base did not change between 1-10 s (Figure 1M; the neck width; 1 s, 71.7 ± 3.2 nm, n=42; 10 s, 65.0 ± 3.7 nm, n=31, p > 0.999). These results indicate that Dyn1 is likely necessary for constricting the neck of endocytic pits during ultrafast endocytosis. Serial reconstructions showed no connections from these pits to other structures such as clathrin-coated vesicles or endosomes (Figure S1D), suggesting that these are not bulk membrane invaginations, such as those previously observed following high-frequency stimulation or high potassium application (Raimondi et al., 2011; Wu et al., 2014). These phenotypes were not rescued by the overexpression of Dyn2 (Fig S2). These data suggest that ultrafast endocytosis is likely mediated by Dyn1.

### The splice variant Dyn1xA, but not xB, mediates ultrafast endocytosis

DNM1 is alternatively spliced at the C-terminus after the proline-rich domain (Figure S3A, S3B). Two of these isoforms, xA and xB, are predominantly expressed in neurons (Chan et al., 2010). They share the same phospho-regulated interaction domain (S774 and S778), which binds Syndapin 1 when dephosphorylated (Anggono et al., 2006), but differ at the very end of the C-terminus after residue S845: xA has a highly disordered 20 amino acid extension that provides two additional SH3 binding motifs (Figure S3B, PSRP and PPRP), whereas xB is shorter but contains a calcineurin-binding domain (PxIxIT, Figure S3B). Previous studies suggest that Dyn1xB is specifically involved in bulk membrane uptake after intense neuronal activity (activity-dependent bulk endocytosis) (Cheung and Cousin, 2019; Xue et al., 2011). To confirm that Dyn1 is involved in ultrafast endocytosis, we performed the rescue experiments in *Dyn1,3* DKO by expressing these two isoforms. *Dyn1,3* DKO phenotypes were completely rescued by the overexpression of Dyn1xA in *Dyn1,3* DKO neurons (Figure 1E, 1F,1K-1L). By contrast, the overexpression of Dyn1xB failed to rescue the defect in ultrafast endocytosis (Figure 1G, 1H, 1K-1L). As in *Dyn1,3* DKO, endocytic pits were arrested on the plasma membrane when Dyn1xB was expressed. Interestingly, the stalled pits had a wider opening at the base than those found in *Dyn1,3* DKO (Figure 1M, pit width: DKO 1 s, 71.7 ± 3.2 nm, n=42; Dyn1xB 1 s, 89.2 ± 9.3 nm, n=16; p = 0.60) and were shallower (Figure 1M, pit depth: DKO 1 s, 44.2 ± 6.0 nm, n=42; Dyn1xB 1 s, 30.0 ± 5.7 nm, n=16; p = 0.786). Like the endocytic pits in the DKO neurons, these pits matured over time, but at a much slower rate – the pits became deeper, but the base remained wide (Figure 1M, 1N; pit width: DKO 10 s, 65.0 ± 3.7 nm, n=31; Dyn1xB 10 s, 76.6 ± 9.6 nm, n=21; p = 0.13, pit depth: DKO 10 s, 65.0 ± 7.7 nm, n=31; Dyn1xB 10 s, 53.3 ± 19.3 nm, n=21; p > 0.999), suggesting that Dyn1xB cannot actively participate in ultrafast endocytosis. Consequently, the number of ferritin-positive large endocytic vesicles and endosomes did not increase in these neurons (Figure 1K, 1L). These defects were not due to the phosphorylation status of the shared interaction domain near S774/778 since the expression of the phospho-deficient form (S774/778A) of Dyn1xB did not rescue these phenotypes (Figure 1I, 1J, 1M, 1N). Together these results suggest that the Dyn1xA splice variant is essential for ultrafast endocytosis, and the extended C-terminal domain likely plays a critical role.

### Dyn1xA is associated with the plasma membrane

The ultrastructural data suggest that Dyn1xA mediates ultrafast endocytosis. However, the accumulation of Dyn1xA to endocytic pits may be too slow if it is recruited from the cytosol. To test how Dyn1xA mediates ultrafast endocytosis, we first checked its localization at synapses using confocal fluorescence microscopy. To avoid mislocalization of Dyn1-GFP from its endogenous location due to overexpression, primary hippocampal neurons were transfected with a small amount of the Dyn1-GFP plasmids (500 ng per 125k neurons) and fixed within 20 hours of the transfection in all localization experiments. Cytosolic tdTomato was co-expressed to visualize neurites, and presynapses marked with the Synaptobrevin 2 (Syb2) antibody. Dyn1xA was localized to presynapses and formed distinct puncta in 51.1 ± 2.9 % of these synapses (Figure 2A, 2B). By contrast, Dyn1xB signals were diffuse throughout the axons (Fig. 2C) and only occasionally formed puncta at synapses (Figure 2A, B; 22.4 % of presynaptic terminals). Moreover, Dyn1xB signals were dim within presynaptic terminals and at the remaining puncta (Figure 2C, 2D), indicating that Dyn1xB does not accumulate at presynaptic terminals or efficiently form puncta. Typically, there was only one Dyn1xA punctum per synaptic bouton, but in 10.9 ± 7.4 % of Dyn1xA puncta positive synapses, 2 or more puncta were observed (Figure 2E). These puncta do not represent ongoing endocytic events at synapses since only 1% of synapses exhibit endocytic profiles in resting conditions (Watanabe et al., 2013b, 2014). These puncta were localized near the edge of Syb2 signals (Figure 2F), suggesting that Dyn1xA is localized at the periphery of a synaptic vesicle cluster. Given that ultrafast endocytosis takes place towards the edge of an active zone (Figure S1A) (Watanabe et al., 2013b), this localization pattern indicates that Dyn1xA is likely concentrated near endocytic sites.

**Figure 2.**
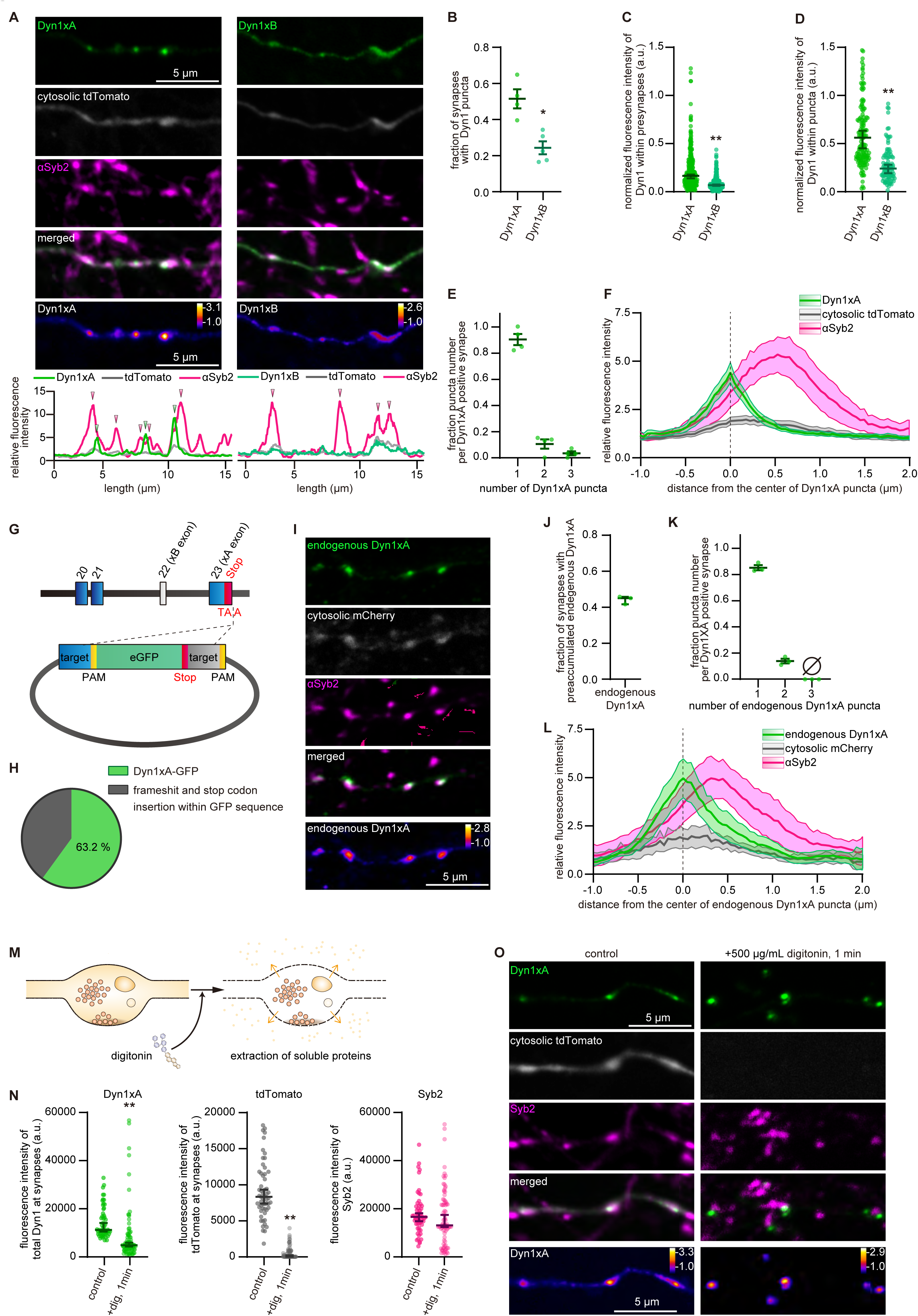
Dyn1xA accumulates within presynapses. (A) Example confocal fluorescence micrographs showing the localization of Dyn1xA-GFP (Dyn1xA) or Dyn1xB-GFP (Dyn1xB), along with exogenously expressed cytosolic tdTomato and antibody staining of Synaptobrevin 2 (*α*Syb2) in fixed neurons. False-colored images of Dyn1xA and Dyn1xB (bottom panels) show the relative fluorescence intensity of these molecules. Line scan graphs represent relative fluorescence intensities of Dyn1xA, Dyn1xB, cytosolic tdTomato, and Syb2 and spatial organizations of these molecules. Green and magenta arrowheads indicating the peaks of Dyn1xA and Syb2 signals, respectively. (B) The fraction of presynapses that contain Dyn1xA or Dyn1xB puncta. n = 4 neurons in Dyn1xA and 5 neurons in Dyn1xB-expressing neurons. To define puncta within synapses, Gaussian smoothing (*σ* = 0.2) was applied on images, and the signals below the average fluorescence intensity of Dyn1xA in the intersynaptic axonal regions removed. The resulting puncta were delineated and set as ROIs. Dyn1xA puncta were defined the puncta adjusting or within Syb2 signals. The median and 95% confidence interval are shown. p = 0.0159, Mann-Whitney test. Each dot represents a fraction calculated from each neuron. (C) The normalized fluorescence intensities of Dyn1xA or Dyn1xB within presynapse. n = 391 Dyn1xA puncta and 350 Dyn1xB puncta. The median and 95% confidence interval are shown. P < 0.0001, Mann-Whitney test. Each dot on the plot represents a punctum. (D) The normalized fluorescence intensities of Dyn1xA or Dyn1xB within puncta. n = 166 Dyn1xA puncta and 96 Dyn1xB puncta. The median and 95% confidence interval are shown. p < 0.0001, Mann-Whitney test. Each dot on the plot represents a punctum. (E) Relative frequency distributions of the number of puncta within presynaptic boutons among those that contain at least one punctum. The fraction is calculated from each neuron. The mean and SEM are shown. n = 4 neurons. (F) The distributions of cytosolic tdTomato and *α*Syb2 relative to the peak of Dyn1xA signals. Line scans of fluorescence from 39 synapses, defined by the end-to-end Syb2 signals, are aligned based on the peak pixel of Dyn1xA signals, and fluorescence intensities averaged. The median and 95% confidence interval are shown. (G) Schematics showing the knock-in strategy for tagging Dyn1xA at the endogenous locus with GFP in primary cultured hippocampal neurons. The gene structure of Dyn1 (top) and the schematic of the knock-in construct (bottom) are shown. eGFP was inserted into the stop codon after the xA exon. (H) The proportion of successful Dyn1xA-GFP knock-in neurons. See Figure S4 for the detail genomic sequencing analysis. (I) Example confocal micrographs showing endogenous localization of Dyn1xA-GFP (endogenous Dyn1xA, stained with anti-GFP antibodies), cytosolic mCherry and *α*Syb2 in fixed neurons. False-colored images representing the relative fluorescence intensity of endogenous Dyn1xA. Cytosolic mCherry was co-overexpressed with the knock-in construct. (J) The fraction of presynapses that contain endogenous Dyn1xA puncta. The mean and SEM are shown. n = 3 neurons. (K) Same as (E), but for endogenous Dyn1xA. The fraction is calculated from each neuron. The mean and SEM are shown. n = 3 neurons. (L) Same as (F), but for endogenous Dyn1xA. The median and 95% confidence interval are shown. n = 14 synapses. (M) Schematics showing the cytosolic extraction using digitonin. (N) Fluorescence intensities of Dyn1xA, cytosolic tdTomato or *α*Syb2 within presynapses in control and digitonin treated neurons. Neurons were fixed at 1 minute after the addition of 500 μg/mL of digitonin and immunostained with anti-Syb2 antibodies. Each dot represents a punctum. The median and 95% confidence interval are shown. n = 57 presynapses (from 4 coverslips) in control and 79 presynapses (from 4 coverslips) in digitonin treated neurons. p < 0.0001 for Dyn1xA and tdTomato in digitonin treated neurons against untreated controls, Mann-Whitney test. (O) Example confocal micrographs showing Dyn1xA, cytosolic tdTomato and *α*Syb2 in control and digitonin-treated neurons after fixation. False-colored images representing the relative fluorescence intensity of Dyn1xA. N = 2 or more independent cultures. *p < 0.05, **p < 0.0001. See Quantification and Statistical Analysis for the n values and detailed numbers.

Since the overexpression of dynamin may result in mislocalization of proteins, we localized endogenous Dyn1xA by inserting GFP at the end of the exon 23 of the DNM1 locus using CRISPR/Cas9 guided gene-editing with Open Resource for the Application of Neuronal Genome Editing (ORANGE) (Willems et al., 2020) (Figure 2G). Sequencing confirmed that GFP was correctly inserted in 63.2 % of neurons (Figure 2H). In the rest, the GFP gene was disrupted by the insertion of a stop codon (Figure 2H, S3C, S3D). Thus, all GFP-positive neurons contain the correct Dyn1xA-GFP sequence in their genome. In these neurons, Dyn1xA puncta were present in 43.5 ± 1.6 % of presynapses (Figure 2I, 2J). Of these, 85.1 ± 1.9 % of Dyn1xA positive synapses contained only one punctum (Figure 2K). Endogenous Dyn1xA puncta were also localized near the edge of Syb2 signals (Figure 2L). These results were nearly identical to the overexpressed exogenous Dyn1xA-GFP signals (Figure 2A, 2F), indicating that our expression scheme can probe the endogenous location of Dyn1xA and that Dyn1xA normally forms puncta within presynaptic terminals, presumably near endocytic sites.

To test whether Dyn1xA puncta are associated with the plasma membrane, we examined the exogenous Dyn1xA-GFP signals while applying digitonin in the external solution (500 µg/ml). Digitonin is a non-ionic weak detergent that makes the plasma membrane porous and permits the diffusion of cytosolic materials from the cell, while retaining organelles inside the cell (Dubreuil et al., 2020; Gopal et al., 2017; Moore and Blobel, 1992) (Figure 2M). Thus, if Dyn1xA puncta were cytosolic, they would diffuse out of cells after the digitonin treatment. As controls, cytosolic tdTomato was co-expressed in these neurons, and synaptic vesicles marked with Syb2 antibodies. Cells were fixed immediately after the treatment. As expected, tdTomato was washed out completely, while synaptic vesicles marked by Syb2 remained within the terminals after the digitonin application for 1 min (Figure 2N, 2O). Although the total fluorescence intensity level decreased by ∼40% (Figure 2N), Dyn1xA puncta remained near the Syb2 signals (Figure 2O), suggesting that these puncta are associated with membranes. Together, these results demonstrate the presence of pre-accumulated Dyn1xA molecules on the plasma membrane at presynaptic terminals.

### Dyn1xA puncta are phase-separated

Dyn1xA forms puncta on the plasma membrane in fixed neurons. To better observe the behavior of Dyn1xA signals, we performed live-cell imaging on neurons expressing Dyn1xA-GFP and Syb2-mCherry (Figure 3A). Immediately after the addition of digitonin, Dyn1xA signals outside the puncta disappeared (Figure 3A-C), indicating that there is a cytosolic pool of Dyn1xA present at presynaptic terminals. By contrast, the puncta remained intact over the course of the experiments (1 min), although the fluorescence intensity gradually decreased to ∼61.9% (Figure 3B, 3C), suggesting that Dyn1xA molecules can slowly diffuse out of the puncta. Furthermore, these puncta were roughly circular with an aspect ratio (AR; maximal diameter/minimal diameter) of 1.23 (Figure 3D) and occasionally underwent fusion and fission (Figure 3E). By contrast, synaptic vesicle clusters did not exhibit such shapes (Figure 3D: AR = 1.68, P < 0.0001). These data are consistent with a liquid droplet state of molecules (Hyman and Brangwynne, 2011; Hyman et al., 2014; Stone, 1994) and suggest that Dyn1xA may be phase-separated.

**Figure 3.**
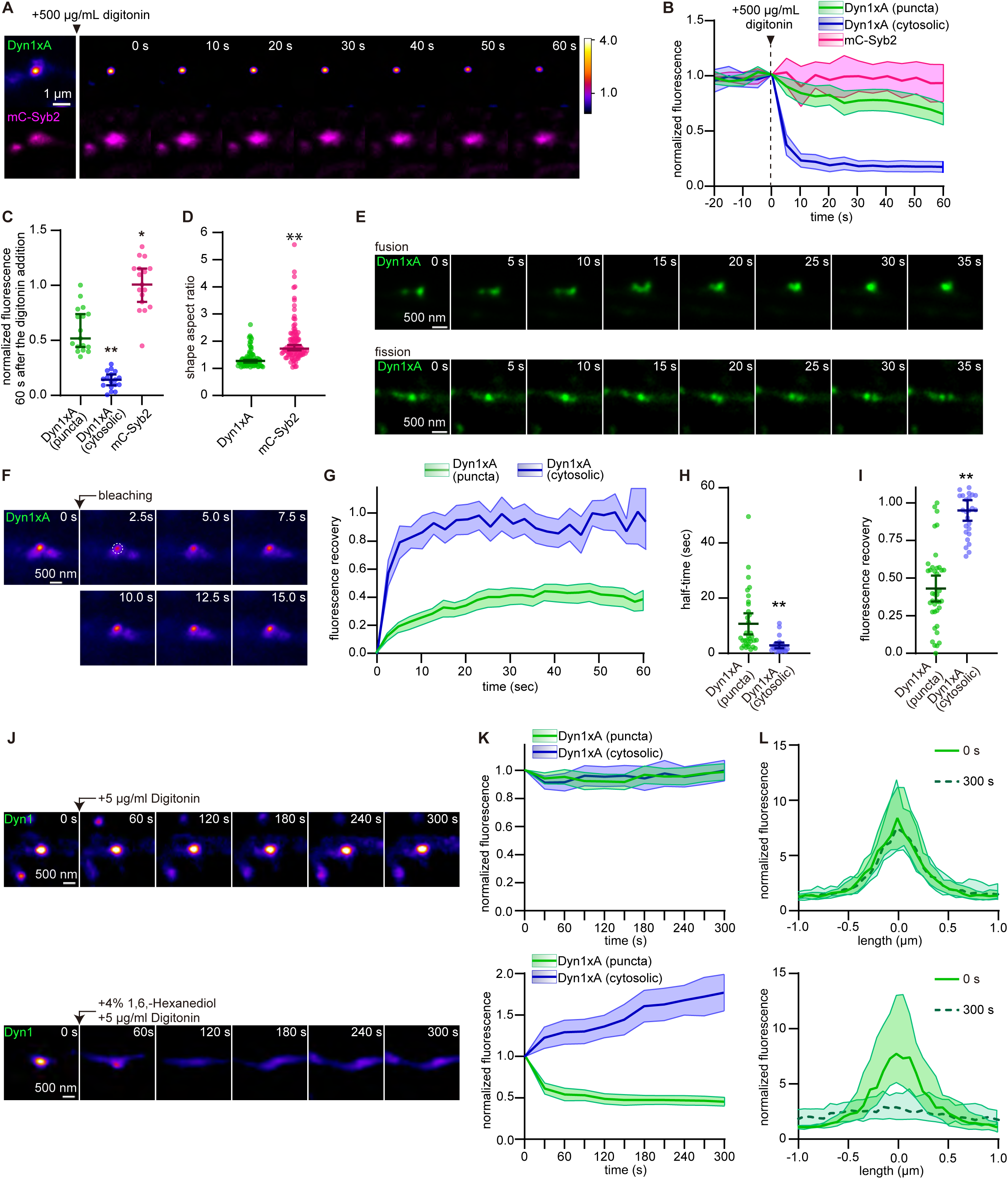
Dyn1xA puncta exhibit liquid-like behaviors. (A) Example live-cell fluorescence micrographs showing overexpression of Dyn1xA and mCherry-Syb2 (mC-Syb2) before and after the digitonin treatment. Time indicates after the 500 μg/mL of digitonin addition. Images are false-colored to indicate relative intensity. (B) Averaged normalized fluorescence intensities of Dyn1xA puncta, cytosolic Dyn1xA and mC-Syb2 before and after the digitonin addition along the time course. Fluorescence is normalized to the pre-treatment. To define puncta, Gaussian smoothing (*σ* = 0.2) was applied on images, and the signals below the average fluorescence intensity of Dyn1xA or Syb2 in the intersynaptic axonal regions removed. The resulting puncta that appeared adjacent to or within Syb2 signals were delineated and set as ROIs. Any signals outside the Dyn1xA ROIs and around Syb2 signals are defined as cytosolic Dyn1xA. The median and 95% confidence interval are shown. (C) Fluorescence intensities of Dyn1xA puncta, cytosolic Dyn1xA within presynaptic boutons, and mC-Syb2, re-plotted from the 60 s time point in (B). The median and 95% confidence interval are shown in each graph. p = 0.0008 for cytosolic Dyn1xA and 0.0188 for mC-Syb2 against Dyn1xApuncta. Kruskal–Wallis tests with comparisons against Dyn1xA by post hoc Dunn’s multiple comparisons tests. For (A-C), 17 different presynapses in 3 different neurons were examined from 2 independent cultures. (D) The aspect ratio (maximum diameter/minimum diameter) of Dyn1xA puncta and mC-Syb2 in live neurons. The aspect ratio of 1 indicates a perfect circle. Each dot on the plot is a punctum. N = 82 Dyn1xA puncta, and n = 85 mC-Syb2 puncta. P< 0.0001. 5 different neurons were examined from 2 independent cultures. (E) Example live cell images showing Dyn1xA puncta undergoing fusion (top) and fission (bottom) events. Times after the initiation of image acquisitions are indicated. (F) Example live-cell fluorescence images showing Dyn1 puncta pre-and post-photobleaching. Times after the photobleaching are indicated. A dashed circle indicates the photobleached region. (G) Fluorescence recovery of Dyn1xA signals when signals at the entire puncta or cytosolic Dyn1xA in axons were photobleached. Times indicate after the photobleaching. The median and 95% confidence interval are shown. (H) The recovery half-time of Dyn1xA signals following the photobleaching of the entire puncta or cytosolic Dyn1xA in axons. The half-time is calculated at the recovery period of 60 s after photobleaching. The median and 95% confidence interval are shown. Each dot represents a punctum. p < 0.0001 for cytosolic Dyn1xA against Dyn1xA puncta. Mann-Whitney test. (I) The fraction of fluorescence recovery 60 s after photobleaching the entire puncta and cytosolic Dyn1xA in axons. The median and 95% confidence interval are shown. Each dot represents a punctum. p <0.0001. Mann-Whitney test. n = Dyn1xA puncta, 71; and cytosolic Dyn1xA, 26; in (G-I). 5 different neurons were examined in each condition from 2 independent cultures. (J) Example live-cell fluorescence micrographs showing Dyn1xA signals over 5 min with the addition of either 5μg/mL digitonin alone (control) or 5μg/mL digitonin + 4% 1,6-hexanediol (1, 6,-hexanediol treatment). Mild concentration of digitonin (5μg/mL) was added to permeabilize the plasma membrane to infuse 1,6-hexanediol into the cells. Times after the addition are shown. (K) Averaged normalized fluorescence intensities of Dyn1xA puncta and cytosolic Dyn1xA in control or 1, 6, -hexanediol treatment. Fluorescence is normalized to the pre-treatment (0 s). Time indicates after the addition. The median and 95% confidence interval are shown. (L) Example line scan normalized fluorescence intensity plot of Dyn1xA around presynaptic terminals in control or 1, 6, -hexanediol treatment. The peak of Dyn1xA fluorescence is centered in the plot (0 µm). Solid line indicates before addition of 5 μg/mL digitonin (top) or the addition of 5 μg/mL digitonin and 4% 1, 6, -hexanediol (bottom). Dashed line indicates 300 s after the treatment. Fluorescence intensities are normalized between the average fluorescence intensities of −1.0 to −0.7 region in control. The median and 95% confidence interval are shown. n = 42 presynapses for the control and 40 presynapses for the 1, 6, -hexanediol treatment in (J-L). 6 different neurons for the control and 4 neurons for the 1, 6, -hexanediol treatment were examined in each condition from 2 independent cultures.

To test whether Dyn1xA molecules form liquid condensates, we first performed fluorescence recovery after photobleaching (FRAP) experiments in neurons expressing Dyn1xA-GFP. When we photo-bleached the Dyn1xA signals at intersynaptic regions along the axon, the signals recovered nearly 100% (100.3 % with a median time constant of 2.1 s) (Figure 3F-3I). The diffusion coefficient was 0.42 µm^2^/s, which is within typical diffusion rates of proteins in the cytosol (Kwapiszewska et al., 2019; Lommerse et al., 2005), indicating that the signals outside the puncta are likely cytosolic and freely diffusing. Interestingly, when the entire Dyn1xA puncta were photobleached, the recovery of fluorescence signals was much slower with a median time constant of 6.3 s (Figure 3G), the fluorescence intensity only recovered to 39.0 % (Figure 3I) and the diffusion coefficient was 0.06 µm^2^/s, suggesting that the diffusion of molecules into the puncta is much slower. These results indicate that Dyn1xA puncta are compartmentalized and form phase boundaries from their surroundings, similarly to what has been described in many biological systems (Brangwynne et al., 2009; Hyman et al., 2014).

A liquid phase of proteins is generated through weak hydrophobic interactions. Previous studies demonstrate that 1,6-hexanediol, an aliphatic alcohol, can disrupt such interactions and dissolve a variety of liquid droplets including stress granules, P bodies, and endocytic proteins (Gopal et al., 2017; Kroschwald et al., 2015; Patel et al., 2007; Kozak and Kaksonen, 2019; Wilfling et al., 2020). Thus, if Dyn1xA forms liquid droplets, the puncta should be dissolved with the application of 1,6-hexanediol. Since 1,6-hexanediol is not membrane-permeable, we used a mild concentration of digitonin (5 µg/ml) to let it into cells. In control neurons treated with only digitonin, the puncta were stable over 5 min (Figure 3J-3L). However, when we added 1,6-hexanediol with digitonin, Dyn1xA puncta were completely dissolved and became diffuse along the axon within 1-2 min (Figure 3J-3L). These data suggest that Dyn1xA puncta are phase-separated on the plasma membrane and hydrophobic protein-protein interactions are necessary for the formation of the pre-accumulated Dyn1xA puncta.

### The phase separation of Dyn1xA requires dephosphorylation of the PRD

The phase separation of proteins requires intrinsically-disordered regions as well as multivalent interactions with other proteins, and is often controlled by posttranslational modifications (Hyman et al., 2014; Milovanovic et al., 2018). The proline-rich domain of Dyn1 satisfies all these requirements. The PRD is the only highly disordered region of Dyn1 (Figure S3A), and the extended PRD of Dyn1xA may provide better interaction sites for multiple endocytic proteins than that of Dyn1xB. To test this possibility, we first performed pull-down assays from mouse nerve terminal lysates using recombinant GST-Dyn1xA-PRD (751-864) and GST-Dyn1xB-PRD (751-851) as baits. We blotted for Syndapin 1, a brain-specific isoform of syndapin known to bind Dyn1 (Anggono et al., 2006) (Figure S4A-4D), and two isoforms of Endophilin A (Endophilin A1 and Endophilin A2) (Figure S4E-4L) involved in ultrafast endocytosis (Watanabe et al., 2018). Syndapin 1 bound equally well to both Dyn1xA and xB (Figure S4D). In contrast, Endophilin A seemed to show a preference: Endophilin A1 but not A2 bind better with Dyn1xA than Dyn1xB (Figure S4H, S4L, Endophilin A1, 1.8-fold: and Endophilin A2, 0.9-fold). Interestingly, the interaction of Dyn1xA with Syndapin 1 was strongly regulated by the phosphorylation status of the PRD: Syndapin 1 binding was reduced by ∼70 % when the Dyn1xA PRD was mutated to be phosphomimetic at S774/778E, but not S851/857E (Figure S4A, S4B). There was no further reduction in binding when S851/857E are additionally introduced to Dyn1xA-PRD S774/778E (Figure S4A, S4B), suggesting that Syndapin 1 binding is strongly regulated at S774/778 of Dyn1xA PRD, as previously reported (Anggono et al., 2006). In contrast, Endophilin A binding to Dyn1xA PRD did not show strong dependence on the phosphorylation status, as previously reported (Anggono et al., 2006) (except for Endophilin A1 to S774/778E, ∼60 % reduction; Figure S4E, 4F, 4I, 4J). These data suggest that Dyn1xA likely allows multivalent interactions with both Syndapin 1 and Endophilin A, and the Syndapin 1 binding is strongly regulated by the phosphorylation status of the PRD.

To determine if phase separation depends on dephosphorylation, we performed photo-bleaching experiments in neurons expressing Dyn1xA-GFP. We applied calcineurin inhibitor FK506 (2 µM, 30 min) to block further dephosphorylation of Dyn1xA or GSK3b inhibitor (CHIR990201, 10 µM, 30 min) to block the phosphorylation of S774. DMSO (0.05%) was applied in controls. Consistent with the results in wild-type neurons (Figure 3G, 3I), Dyn1xA-GFP signals recovered to median 39.0% within 1 min of photobleaching in the DMSO-treated cells (Figure S5A-C). The FK506 treatment had no effect on recovery (Figure S5A-C), presumably because ∼75% of Dyn1xA is not phosphorylated in the resting condition (Anggono et al., 2006; Liu et al., 1994). By contrast, the level of fluorescence recovery was enhanced in CHIR990201-treated neurons (median 59.1 %) (Figure S5A-S5C). These data indicate that dephosphorylation increases the amount of cytosolic Dyn1xA that can be exchanged with those in puncta.

To further examine the requirement for the dephosphorylation of Dyn1xA in phase separation, we localized wild-type, phosphomimetic (S774/778D) and phospho-deficient (S774/778A) forms of Dyn1xA-GFP in wild-type neurons. Cytosolic tdTomato was co-expressed, and synaptic vesicle clusters visualized with Syb2 antibodies. As in wild-type Dyn1xA-GFP (Figure 2A and 2F), a phospho-deficient Dyn1xA-GFP appeared punctate along the axons and localized at the edge of the synaptic vesicle clusters (Figure 4A). More than a half of putative synapses (58.2% of Syb2-positive boutons) contained at least one punctum when a phospho-deficient form was expressed, similarly to when a wild-type form was expressed (65.1%, p > 0.999) (Figure 4A, 4B). The total fluorescence levels of Dyn1xA in boutons were similar between neurons expressing wild-type and phospho-deficient Dyn1xA-GFP (P>0.9999, Figure 4C), but fluorescence signals within puncta were brighter by 25.8% in neurons expressing the phospho-deficient form (Figure S6), suggesting that phosphodeficient Dyn1xA accumulate in the puncta efficiently. By contrast, the phosphomimetic form appeared diffuse throughout the axons; only 26.9% of Syb2-positive boutons contained puncta (Figure 4A, 4B, p = 0.02). The total fluorescence level of Dyn1xA within boutons was reduced by 44.5% (P<0.0001, Figure 4C), and the remaining puncta in these boutons were dim: the fluorescence intensity was reduced by 42% (P<0.0001, Figure S6), suggesting that Dyn1xA molecules do not accumulate in the puncta when Dyn1xA is phosphorylated. Together, these results suggest that Dyn1xA forms liquid condensates on the plasma membrane at synapses, and this formation is promoted by dephosphorylation.

**Figure 4.**
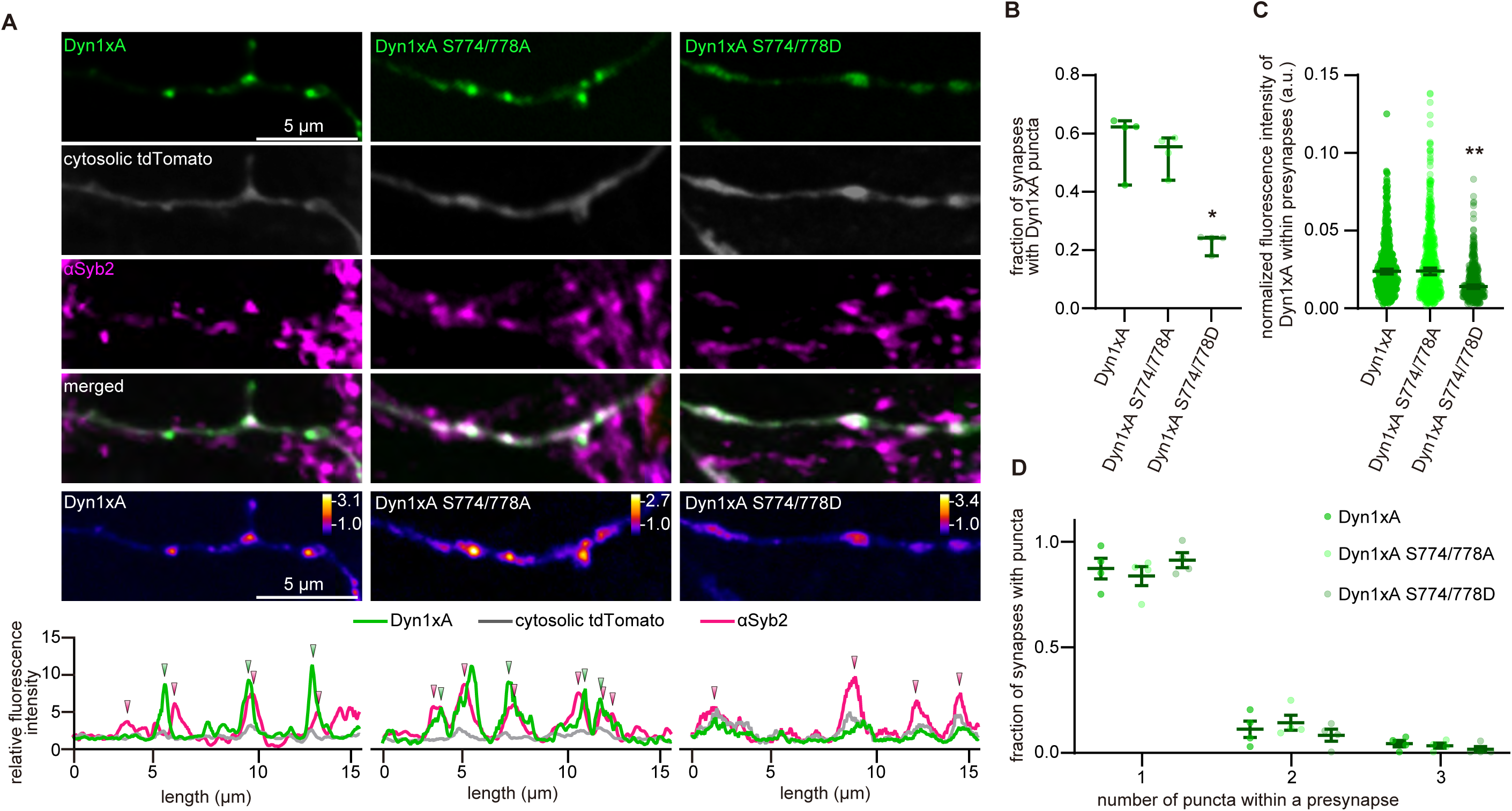
Dyn1xA phase separation requires dephosphorylation of the proline-rich domain. (A) Example confocal micrographs showing overexpression of Dyn1xA, Dyn1xA S774/778A-GFP (Dyn1xA S774/778A) or Dyn1xA S774/778D-GFP (Dyn1xA S774/778D), along with exogenously expressed cytosolic tdTomato and immuno-stained *α*Syb2 in fixed neurons. False-colored images (bottom panels) show the relative fluorescence intensity of Dyn1xA, Dyn1xA S774/778A or Dyn1xA S774/778D. Line scan graphs represent the relative locations of Dyn1xA, Dyn1xA S774/778A or Dyn1xA S774/778D to cytosolic tdTomato and *α*Syb2. (B) The fraction of presynapses that contain Dyn1xA, Dyn1xA S774/778A or Dyn1xA S774/778D puncta. n = 4 neurons in each case. The median and 95% confidence interval are shown. p > 0.999 for Dyn1xA S774/778A and = 0.0181 for Dyn1xA S774/778D against Dyn1xA. Kruskal–Wallis tests with full pair-wise comparisons against Dyn1xA by post hoc Dunn’s multiple comparisons tests. (C) The normalized fluorescence intensities of Dyn1xA, Dyn1xA S774/778A or Dyn1xA S774/778D within presynaptic terminals. The median and 95% confidence interval are shown. Each dot on the plot is a presynapse. n = 408 Dyn1xA, 571 for Dyn1xA S774/778A, and 444 Dyn1xA S774/778D presynapses. The median and 95% confidence interval are shown. p > 0.999 for Dyn1xA S774/778A and < 0.0001 for Dyn1xA S774/778D against Dyn1xA. Kruskal–Wallis tests with full-pair wise comparisons against Dyn1xA by post hoc Dunn’s multiple comparisons tests. (D) Relative frequency distributions of the number of puncta within presynaptic boutons among those that contain at least one punctum in neurons expressing Dyn1xA, Dyn1xA S774/778A or Dyn1xA S774/778D puncta within presynapse. The mean and SEM are shown. The fraction is calculated from each neuron. The mean and SEM are shown. n = 4 neurons in each case. N = 2 independent cultures. *p < 0.05, **p < 0.0001. See Quantification and Statistical Analysis for the n values and detailed numbers.

### Enhanced synaptic activity increases the number of Dyn1xA puncta

When the phospho-deficient form of Dyn1xA was expressed, there was an increase in the fluorescence intensity within puncta (Figure 4D), suggesting the amount of Dyn1xA within puncta is controlled by phosphorylation status. Since the dephosphorylation of Dyn1 is dependent on the synaptic activity level (Anggono et al., 2006), we applied 90 mM KCl (90 s) to induce the depolarization of the plasma membrane. Physiological saline solution with 2.5 mM KCl (sham) was applied in control neurons (Figure S7). The fraction of putative synapses with puncta did not change under these treatments (Figure S7B). However, when high K+ is applied to neurons, the fluorescence levels of Dyn1xA within synapses and puncta increased significantly (Figure S7C, S7D). The diameter of these puncta was substantially larger than that of puncta after 2.5 mM K+ (Figure S7E). Furthermore, synapses that contain 2 puncta increased, while those that contain 1 punctum decreased (Figure S7F), suggesting that more Dyn1xA becomes available when the synaptic activity level increases. These data indicate that the number of Dyn1xA puncta is modulable commensurate to activity levels of synapses.

### Dephosphorylated Dyn1xA is required for the kinetics of ultrafast endocytosis

The phosphorylation status of Dyn1xA controls its membrane localization and phase separation. This membrane localization would likely accelerate endocytosis by bypassing the slow recruitment of molecules to endocytic sites. Since the phospho-deficient form of Dyn1xA is associated with the plasma membrane similarly to wild-type Dyn1xA (Figure 1), ultrafast endocytosis is expected to take place normally when this form is expressed. By contrast, the phosphomimetic form is largely cytosolic. Having to recruit these molecules from the cytosol would likely slow down ultrafast endocytosis. To test these possibilities, we performed ‘flash-and-freeze’ experiments in *Dyn1,3* DKO neurons, expressing wild-type, phospho-deficient (S774/778A), or phosphomimetic (S774/778D) forms of Dyn1xA (Figure 5). The same conditions as in Figure 1 were used (single stimulus, 37 °C, 4 mM Ca^2+^, TTX incubation overnight, and ferritin incubation for 5 min). *Dyn1,3* DKO and *Dyn3* KO were used as additional controls. The results from the controls were as expected: *Dyn3*KO showed normal ultrafast endocytosis, while *Dyn1,3* DKO exhibited stalled endocytic pits on the plasma membrane (Figure 5A-5D, 5K, 5L). The endocytic defect of *Dyn1,3* DKO was rescued by the phospho-deficient and wild-type forms. Ferritin particles were found in endocytic vesicles and endosomes by 1 s and additionally in synaptic vesicles by 10 s (Figure 5E-5G, 5K, 5L). However, in neurons expressing the phosphomimetic form, endocytic pits were arrested at the plasma membrane up to 10 s after stimulation (Figure 5I-5L). Consequently, the number of ferritin-positive vesicles and endosomes did not increase over time, suggesting that the phosphomimetic form cannot rescue the ultrafast endocytic defect. Interestingly, unlike endocytic pits in *Dyn1,3* DKO, which exhibited a wide opening at their base (Figure 5M; pit width, 1s, median 61.67 nm, n=57 pits; 10 s, median 58.33 nm, n=50 pits, p > 0.999), the base of endocytic pits in Dyn1xA S774/778D neurons was constricted over time and formed a neck (Figure 5I, 5M, 5N; pit width, 1 s: pit width, median 58.33 nm, n=34 pits; 10 s, median 33.33 nm, n=43 pits, P<0.0001), suggesting that endocytosis may complete in the presence of Dyn1xA S774/778D but at a much slower rate. These results are consistent with previous studies using pHluorin assays (Armbruster et al., 2013), indicating that these phosphorylation sites control the kinetics of endocytosis. Given that the majority of Dyn1xA S774/778D is cytosolic and diffuse throughout axons (Figure 4A), ultrafast endocytosis is likely delayed due to the slow recruitment of these molecules to endocytic sites. Altogether, these data suggest that the phase separation of Dyn1xA accelerates endocytosis at synapses by bypassing the recruitment step of endocytosis.

**Figure 5.**
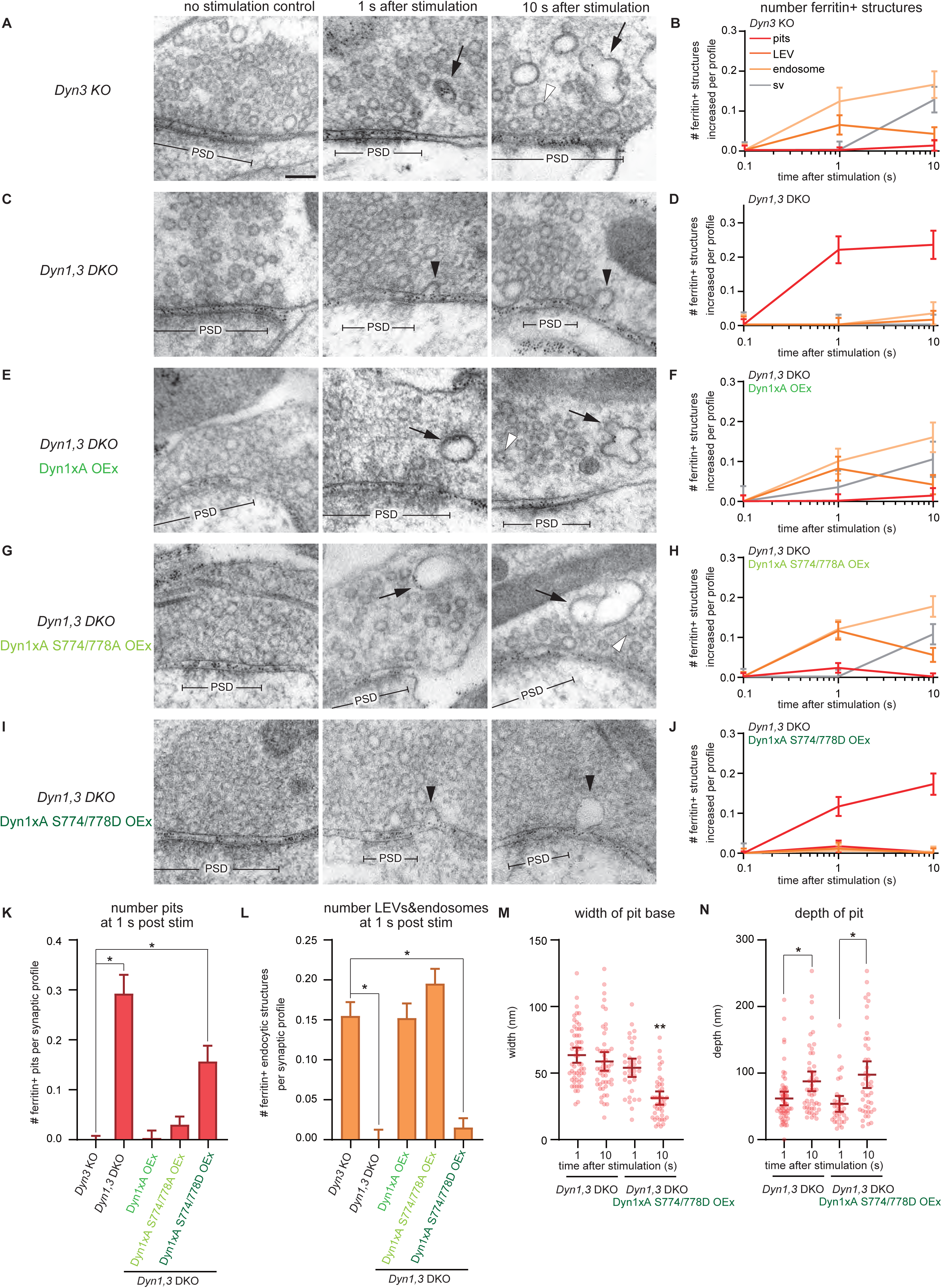
Dephosphorylation of Dyn1xA is required for the kinetics of ultrafast endocytosis. (A, C, E, G and I) Example micrographs showing endocytic pits and ferritin-containing endocytic structures at the indicated time points in *Dyn3* KO (A), *Dyn1,3* DKO (C), *Dyn1,3* DKO, Dyn1xA overexpression (OEx) (E), *Dyn1,3* DKO, Dyn1xA S774/778A OEx (G), *Dyn1,3* DKO, Dyn1xA S774/778D OEx (I). Black arrowheads, endocytic invaginations; black arrows, LEVs or endosomes; white arrowheads, synaptic vesicles. Note that the endocytic defect can be rescued with Dyn1xA S774/778A but not with Dyn1xA S774/778D. Scale bar: 100 nm. PSD, post-synaptic density. (B, D, F, H and J) Plots showing the increase in the number of each ferritin-positive endocytic structure per synaptic profile after a single stimulus in *Dyn3* KO (B), *Dyn1, 3* DKO (D), *Dyn1, 3* DKO, Dyn1 OEx (F), *Dyn1, 3* DKO, Dyn1xA S774/778A OEx (H) and *Dyn1, 3* DKO, Dyn1zA S774/778D OEx (J). The mean and SEM are shown in each graph. (K) Number of endocytic invaginations at 1s after the stimulations. The numbers are re-plotted as a bar graph from the 1 s time point in (B, D, F, H, and J) for easier comparison between groups. P = 0.0037 for *Dyn1,3* DKO and 0.0366 for *Dyn1,3* DKO, Dyn1xA S774/778D OEx against *Dyn3* KO. Ordinary one-way ANOVA with full pairwise comparisons by Holm-Šídák’s multiple comparisons test. The mean and SEM are shown. (L) Number of LEVs and endosomes at 1s after stimulation. The numbers of LEVs and endosomes are summed from the data presented in (B, D, F, H, and J), averaged, and re-plotted for easier comparison between groups. p = 0.0079 for *Dyn1,3* DKO and 0.0093 for *Dyn1,3* DKO, Dyn1xA S774/778D OEx against *Dyn3* KO. Ordinary one-way ANOVA with full pairwise comparisons by Holm-Šídák’s multiple comparisons test. The mean and SEM are shown. (M and N) Plots showing the width (M) and depth (N) of endocytic pits at the 1s time point. The median and 95% confidence interval are shown in each graph. n = *Dyn1, 3* DKO, 57 pits for 1 s and 50 pits for 10 s; *Dyn1, 3* DKO, Dyn1xA S774/778D OEx, 34 pits for 1 s and 43 pits for 10 s. The width; p < 0.0001 for Dyn1xA S774/778D OEx at 10 s against *Dyn1, 3* DKO at 1 s, 10 s and Dyn1xA S774/778D OEx at 1 s. The depth; p = 0.0242 for *Dyn1, 3* DKO for 10 s against *Dyn1, 3* DKO for 1 s; p = 0.0016 for Dyn1xA S774/778D OEx for 10 s against Dyn1xA S774/778D OEx for 1 s. Kruskal-Wallis Test with full comparisons by post hoc Dunn’s multiple comparisons tests. All data are from two independent cultures prepared and frozen on different days. n = *Dyn* 3KO, 693; *Dyn1, 3* DKO, 697; *Dyn1, 3* DKO, Dyn1xA OEx, 655; *Dyn1, 3* DKO, Dyn1xA S774/778A OEx, 728; and *Dyn1, 3* DKO, Dyn1xA S774/778D OEx, 644 synaptic profiles in (B, D, F, H, J, M, and N). See Quantification and Statistical Analysis for the n values and detailed numbers for each time point. Knock out neurons are from the littermates in all cases. * p < 0.05, ** p < 0.0001.

### Syndapin 1 acts as an adaptor between the plasma membrane and Dyn1xA

How does Dyn1xA produce molecular condensates on the plasma membrane? Dyn1xA has a PH domain (Figure S3A, S3B) that interacts with phosphatidylinositol phosphates (PIPs) and can bind membranes by itself. However, the experiments with the phosphomimetic form of Dyn1xA suggest an essential function of the PRD in the phase separation. Since the Dyn1xA PRD interacts with the SH3 domain of Syndapin 1 (Figure S4) and their interaction is strongly controlled by the phosphorylation status of S774/S778 (Figure S4B) (Anggono et al., 2006), we tested whether Syndapin 1 has a role in the phase separation of Dyn1xA (Figure 6). We generated shRNA against Syndapin 1 (Syndapin 1 knock-down or KD, hereafter) and infected neurons on DIV3 with a lentivirus carrying this vector. The knock-down efficiency was ∼80% (Figure S8, DIV14-15). Scramble shRNA was used as a control (Figure S8). The Dyn1xA-GFP and cytosolic tdTomato constructs were transfected ∼20 hours before the experiments, and neurons fixed just prior to imaging. Presynaptic terminals were visualized using Syb2 antibodies. In scramble shRNA controls, Dyn1xA-GFP formed puncta along the axons and localized next to Syb2 signals (Figure 6A, 6B; 53.2% of Syb2-positive boutons) – consistent with the results in the wild type (Figure 2). However, in Syndapin 1 KD, Dyn1xA-GFP signals were diffuse along axons and only occasionally formed puncta (Figure 6A-6E; 20.3% of Syb2-positive boutons), suggesting that Syndapin 1 is necessary for the localization of Dyn1xA. The total fluorescence level of Dyn1xA in boutons was reduced (Figure 6C; 66.6 % reduction in normalized fluorescence intensity). The remaining puncta were dim (Figure 6D; 45.3 % reduction in normalized fluorescence intensity). Distribution of puncta number per bouton did not change between scramble shRNA and Syndapin 1 shRNA (Figure 6E). These phenotypes were similar to those observed in neurons expressing Dyn1xA S774/778D-GFP, suggesting that Syndapin 1 is an essential interaction partner of Dyn1xA for the formation of liquid-like molecular condensates.

**Figure 6.**
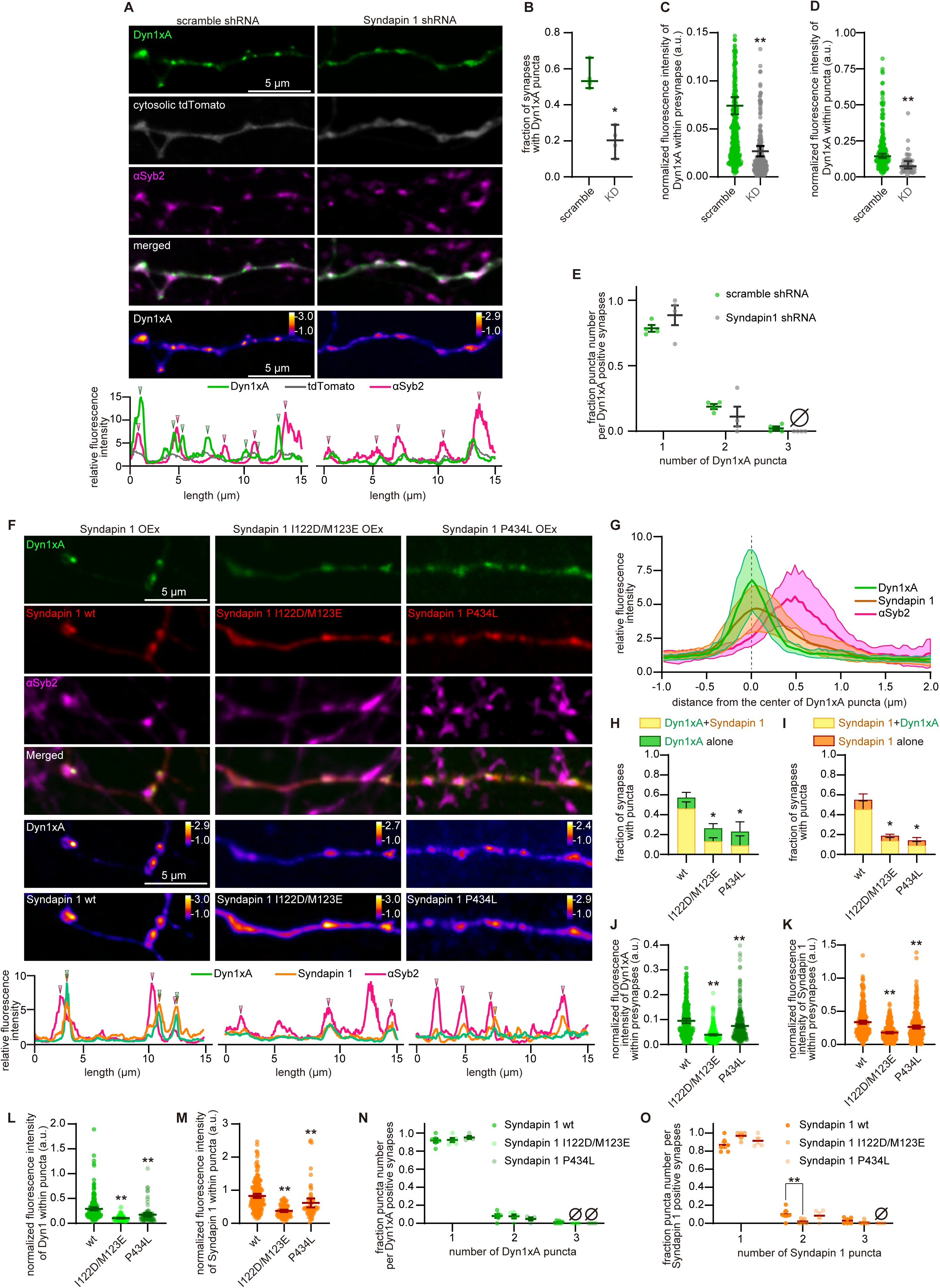
Syndapin 1 is essential for the phase separation of Dyn1xA. (A) Example confocal immunofluorescence micrographs showing overexpression of Dyn1xA, cytosolic tdTomato, and *α*Syb2 in neurons expressing scramble shRNA or Syndapin 1 shRNA. False-colored images (the bottom panels) show the relative fluorescence intensity of Dyn1xA. Line scan graphs represent the localization of Dyn1xA relative to cytosolic tdTomato and *α*Syb2. (B) The fraction of presynapses that contain Dyn1xA in neurons expressing scramble shRNA or Syndapin 1 shRNA. n = 4 neurons in both scramble and Syndapin 1 shRNA expressing neurons. The median and 95% confidence interval are shown. p = 0.0286, Mann-Whitney test. (C) The normalized fluorescence intensities of Dyn1xA within the presynapse in neurons expressing scramble shRNA or Syndapin 1 shRNA. Each dot represents a punctum. n = 388 for scramble and 236 for Syndapin 1 shRNA. The median and 95% confidence interval are shown. P < 0.0001, Mann-Whitney test. (D) The normalized fluorescence intensities of Dyn1xA within the puncta in neurons expressing scramble shRNA or Syndapin 1 shRNA. Each dot represents a punctum. n = 242 for scramble and 44 for Syndapin 1 shRNA. The median and 95% confidence interval are shown. p < 0.0001, Mann-Whitney test. (E) Relative frequency distributions of the number of puncta within presynaptic boutons among those that contain at least one punctum in neurons expressing scramble shRNA or Syndapin 1 shRNA. The mean and SEM are shown in each graph. The fraction is calculated from each neuron. The mean and SEM are shown. n = 4 neurons for both. (F) Example confocal immunofluorescence micrographs showing overexpression of Dyn1xA along with mCherry-Syndapin 1 wild type (Syndapin 1 wt), mCherry-Syndapin 1 I122D/123ME (Syndapin 1 ID/ME), or mCherry-Syndapin 1 P434L (Syndapin 1 P434L). The endogenous Syndapin 1 is knocked-down by shRNA, and all mCherry-Syndapin 1 constructs were rendered shRNA-resistant. Syb2 is immuno-stained to visualize synaptic vesicle clusters (*α*Syb2). Line scan graphs show colocalization of Dyn1xA with *α*Syb2 and wild-type or mutant Syndapin 1 (wt, I122D/M123E, and P434L). (G) The distribution of *α*Syb2 and wild-type Syndapin 1 relative to the peak of Dyn1xA signals. Line scans of fluorescence from 12 synapses, defined by the end-to-end Syb2 signals, were aligned based on the peak pixel of Dyn1xA signals, and fluorescence intensities averaged. The median and 95% confidence interval are shown. (H) The fraction of synapses with Dyn1xA puncta that colocalize (yellow) or does not colocalize with Syndapin 1 (green) wild type (wt) or mutants (I122D/M123E, and P434L). (I) The fraction of synapses with wild type (wt) or mutant (I122D/M123E, and P434L) Syndapin 1 puncta that colocalize (yellow) or does not colocalize with Dyn1xA puncta (orange). (J and K) Normalized fluorescence intensities of Dyn1xA (J) and Syndapin 1 (K) within presynapses of neurons expressing wild-type or mutant (I122D/M123E, and P434L) Syndapin 1. Each dot represents a presynapse. n = 332 for wt, 418 for I122D/M123E, and 341 for P434L. Dyn1xA intensities; p = < 0.0001 for I122D/M123E and 0.0004 for P434L against Syndapin 1 expressing neurons. Syndapin 1 intensities; p = < 0.0001 for I122D/M123E and for P434L against wt expressing neurons. Kruskal–Wallis tests with comparisons against wt by post hoc Dunn’s multiple comparisons tests. The median and 95% confidence interval are shown. (L and M) Normalized fluorescence intensities of Dyn1xA puncta (L) and Syndapin 1 puncta (M) of neurons expressing wild-type or mutant (I122D/M123E and P434L) Syndapin 1. Each dot represents a punctum. Dyn1xA puncta: n = 208 for wt, 120 for I122D/M123E, and 91 for P434L. Syndapin 1 puncta: n = 202 for wt, 88 for I122D/M123E, and 60 for P434L. Dyn1xA intensities; p = < 0.0001 for I122D/M123E and for P434L against Syndapin 1 expressing neurons. Syndapin 1 intensities; p = < 0.0001 for I122D/M123E and for P434L against wt expressing neurons. Kruskal–Wallis tests with comparisons against wt by post hoc Dunn’s multiple comparisons tests. The median and 95% confidence interval are shown. (N and O) Relative frequency distributions of the number of Dyn1xA puncta and Syndapin 1 puncta within presynaptic boutons among those that contain at least one punctum in neurons expressing wild-type or mutant (I122D/M123E and P434L) Syndapin 1. The mean and SEM are shown in each graph. The fraction is calculated from each neuron. The mean and SEM are shown. n = 6 for Syndapin 1 expressing neurons, 5 for I122D/M123E and 4 for P434L. P = 0.0095 for two puncta per synapses in I122D/M123E against Syndapin 1 expressing neurons. Two way ANOVA with Šídák’s multiple comparisons test. In both (A-E) and (F-O) experiments, > 50 boutons were quantified from at least 4 different neurons. N = 2 or more independent cultures. * p < 0.05, ** p < 0.0001. See Quantification and Statistical Analysis for the n values and detailed numbers.

If Syndapin 1 is required for the Dyn1xA phase separation on the membrane, Syndapin 1 should co-localize with Dyn1xA, and mutations in the membrane binding domain (I122/M123) (Rao et al., 2010) or the Dyn1-binding SH3 domain (P434) (Widagdo et al., 2016) should disrupt Dyn1xA localization. To test these predictions, we visualized Dyn1xA-GFP and mCherry-Syndapin 1 localization in the BAR-domain mutant (I122D/M123E) and SH3-domain mutant (P434L). To reduce the overexpression of Syndapin 1, the experiments were performed in Syndapin 1 KD neurons, and the rescue constructs were rendered shRNA-resistant (see Experimental Procedures, Figure S8). The wild-type Syndapin 1 formed puncta along the axons and colocalized with Dyn1xA near the Syb2 signals (Figure 6F, 6G, 6H; 45.0 % of Syb2 positive boutons). By contrast, the Syndapin BAR-domain mutation caused both Syndapin 1 and Dyn1xA to be diffuse along the axons (Figure 6F); colocalization was only observed in 12.4 % of Syb2-positive boutons (Figure 6H, 6I). The total fluorescence levels of Dyn1xA and Syndapin 1 in boutons as well as within puncta were reduced in neurons expressing Syndapin 1-I122D/M123E (Figure 6J-M). Likewise, Dyn1xA and Syndapin 1were diffuse in the axons of neurons expressing the Syndapin SH3 mutant (Figure 6F, 6H, 6I), but more Dyn1xA and Syndapin1 seem to be present at synaptic terminals (Figure 6J-6O). These data suggest that both membrane and protein interacting abilities of Syndapin 1 are necessary for the proper Dyn1xA localization. Thus, Syndapin 1 acts as an adaptor between the plasma membrane and Dyn1xA.

### Syndapin 1 is necessary for the kinetics of ultrafast endocytosis

Syndapin 1 is essential in the pre-recruitment of Dyn1xA on the plasma membrane. When Syndapin 1 is absent or its membrane or protein interacting domains are mutated, Dyn1xA becomes cytosolic. To determine whether the change in Dyn1xA localization due to these mutations has functional consequences, we performed flash-and-freeze experiments in Syndapin 1 KD neurons, expressing shRNA-resistant wild-type, I122D/M123E, or P434L Syndapin 1 (Figure 7, a single stimulus, 37 °C, 4 mM Ca^2+^). Ferritin particles were used as fluid phase markers (2 mg/ml, 5 min). Ultrafast endocytosis and subsequent endosomal sorting were all normal in neurons expressing scrambled shRNA (Figure 7A, 7B). By contrast, Syndapin 1 KD neurons exhibited an accumulation of endocytic pits on the plasma membrane at 1 s (Figure 7C, 7D, 7K, and 7L) – the full resolution of these intermediates required ∼10 s (Figure 7C, 7D, 7K, and 7L), likely because the base of pits was slowly constricted in Syndapin 1 KD (Figure 7M, 7N: pit width, 100 ms, mean 84.7 ± 5.5 nm, n=31 pits; 1 s, mean 77.6 ± 3.7 nm, n=52 pits 10 s, mean 58.5 ± 8.8 nm, n=10 pits, P<0.03 against 100 ms). These defects were rescued by the overexpression of wild-type Syndapin 1, but not with Syndapin BAR domain mutant (I122D/M123E) or SH3 domain mutant (P434L) (Figure 7E-7L). These results were similar to the endocytic defects observed in *Dyn1,3* DKO neurons expressing the phosphomimetic form of Dyn1xA (Figure 5), which cannot interact with Syndapin 1 (Figure S4). Together, these data suggest that the kinetics of ultrafast endocytosis is controlled by the interaction between Syndapin 1 and Dyn1xA and assembly at the plasma membrane before endocytosis.

**Figure 7.**
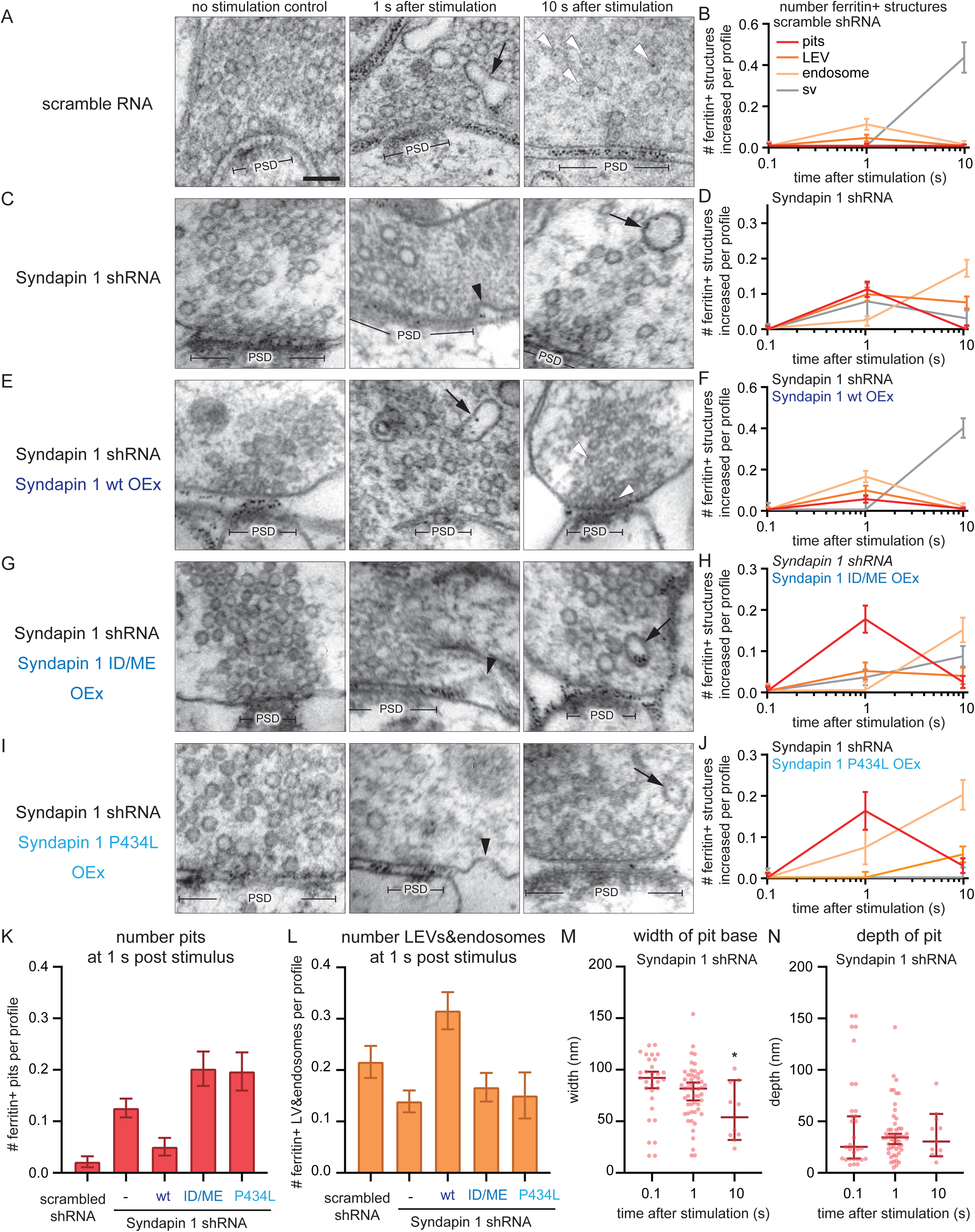
Syndapin 1 is necessary for the kinetics of ultrafast endocytosis. (A, C, E, G and I) Example transmission electron micrographs showing endocytic pits and ferritin-containing endocytic structures at the indicated time points in neurons expressing scramble RNA (A), Syndapin 1 shRNA (C), Syndapin 1 shRNA and wild-type Syndapin 1 overexpression (Syndapin 1 wt OEx) (E), Syndapin 1 shRNA and Syndapin 1 I122D/M123E (Syndapin 1 ID/ME OEx) (G), Syndapin 1 shRNA and Syndapin 1 P434L (Syndapin 1 P434L OEx) (I). In all cases, Syndapin 1 is rendered shRNA-resistant. Black arrowheads, endocytic invaginations; black arrows, ferritin-positive large endocytic vesicles (LEVs) or endosomes; white arrowheads, ferritin-positive synaptic vesicles. Scale bar: 100 nm. PSD, post-synaptic density. (B, D, F, H, and J) Plots showing the increase in the number of each endocytic structure per synaptic profile after a single stimulus in neurons expressing scramble shRNA (B), Syndapin 1 shRNA (D), Syndapin 1 shRNA and Syndapin 1 wt OEx (F), Syndapin 1 shRNA and Syndapin 1 ID/ME OEx (H), Syndapin 1 shRNA and Syndapin 1 P434L OEx (J). The mean and SEM are shown in each graph. (K) Number of endocytic invaginations at 1s after stimulation. The numbers are re-plotted as a bar graph from the 1 s time point in (B, D, F, H, and J) for easier comparison between groups. p = 0.0043 for Syndapin 1 shRNA, <0.0001 for Syndapin 1 shRNA and Syndapin 1 ID/ME OEx and 0.0001 for Syndapin 1 shRNA and Syndapin 1 P434L OEx against scramble RNA. Ordinary one-way ANOVA with full pairwise comparisons by Holm-Šídák’s multiple comparisons test. The mean and SEM are shown. (L) Number of ferritin-positive LEVs and endosomes at 1s after stimulation. The mean and SEM are shown. The numbers of LEVs and endosomes are summed from the data presented in (B, D, F, H, and J), averaged, and re-plotted for easier comparison between groups. p = < 0.0001 for Syndapin 1 shRNA and Syndapin 1 wt OEx, 0.0089 for Syndapin 1 shRNA and Syndapin 1 ID/ME OEx and 0.0093 for Syndapin 1 shRNA and Syndapin 1 P434L OEx against Syndapin 1 shRNA. Ordinary one-way ANOVA with full pairwise comparisons by Holm-Šídák’s multiple comparisons test. The mean and SEM are shown. (M and N) Plots showing the width (M) and depth (N) of endocytic pits at the 1s time point. The median and 95% confidence interval are shown in each graph. n = 31 pits for 0.1 s, 52 pits for 1 s and 10 pits for 10 s. Width; p = 0.290 for 1 s and 0.032 for 10 s against 0.1 s, Kruskal-Wallis test with full pairwise comparisons by post hoc Dunn’s multiple comparisons tests. All data are from at least two independent experiments performed on cultures prepared and frozen on different days. n = scramble RNA, 607; Syndapin 1 shRNA, 1264; Syndapin 1 shRNA and Syndapin 1 wt OEx, 646; Syndapin 1 shRNA and Syndapin 1 ID/ME OEx, 670; and Syndapin 1 shRNA and Syndapin 1 P434L OEx, 603 synaptic profiles in (B, D, F, H, J, M, and N). * p < 0.05, ** p < 0.0001. See Quantification and Statistical Analysis for the n values and detailed numbers for each time point. Knock out neurons are from the littermates in all cases.

## Discussion

### The mode of endocytosis at synapses

A large body of literature suggests that clathrin-mediated endocytosis is responsible for retrieving synaptic vesicle components from the plasma membrane. This conclusion relies on genetic or molecular perturbations that indicate a requirement for clathrin-associated proteins (Dittman and Ryan, 2009; Saheki and De Camilli, 2012). Our data suggest that proteins associated with clathrin-mediated endocytosis also function in clathrin-independent endocytosis. Moreover, these proteins must be clustered at sites of endocytosis to support ultrafast endocytosis. Specifically, we demonstrate that Syndapin I anchors Dynamin 1 in a molecular condensate prior to endocytosis (Figure 8 and Supplementary movie).

**Figure 8.**
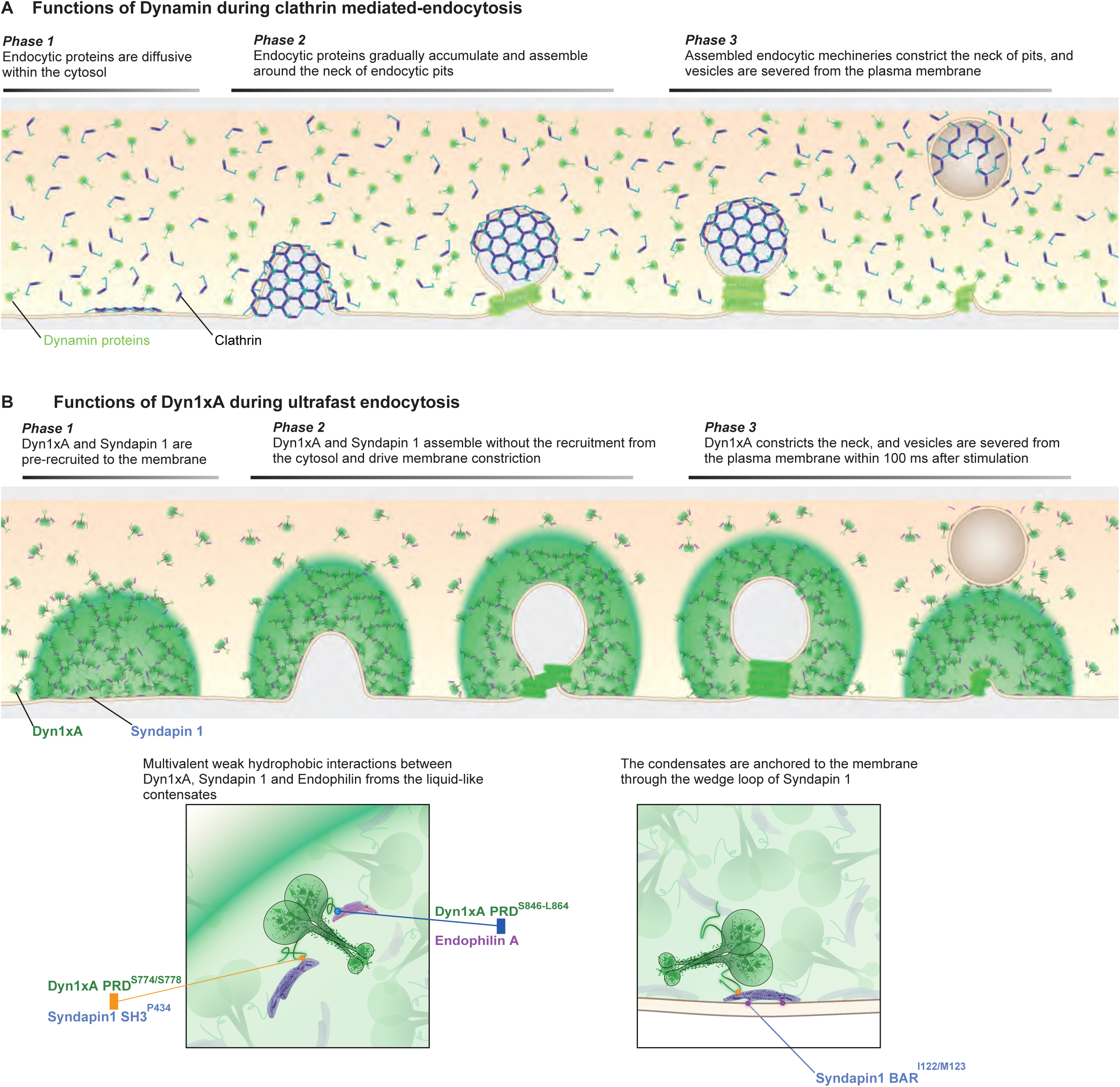
Schematics depicting the differences between clathrin-mediated endocytosis and ultrafast endocytosis. In clathrin-mediated endocytosis (A), dynamin and endocytic proteins are recruited from the cytosol. This process is slow, taking tens of seconds. During ultrafast endocytosis (B), the Dyn1xA splice variant forms phase-separated puncta within presynaptic boutons. The puncta are most likely maintained through the multivalent interaction of Dyn1xA PRD with the SH3 domains of Syndapin 1 and Endophilin A. The puncta are anchored to the plasma membrane, presumably around the active zone, through the wedge loop of Syndapin 1. Loss of interactions among these proteins or the membrane anchor of Syndapin 1 slows down the kinetics of ultrafast endocytosis by 100-fold, since they become cytosolic and need to be recruited to the endocytic sites after the initiation of ultrafast endocytosis.

Other proteins involved in endocytosis are likely to be associated with this matrix. Syndapin also binds Synaptojanin 1 (Qualmann et al., 1999) and Synaptojanin and Dynamin 1 bind Endophilin A (Micheva et al., 1997; Verstreken et al., 2003)(Ringstad et al., 1999). Like Dynamin 1, Endophilin A and Synaptojanin 1 cinch the neck at the base of endocytic pit during ultrafast endocytosis (Watanabe et al., 2018) and are therefore likely to be localized within the Dyn1xA condensate. In the absence of Endophilin A, Synaptojanin 1 becomes diffusely distributed (Milosevic et al., 2011), and consequently, endocytosis slows down substantially when assayed with pHluorin and flash-and-freeze electron microscopy (Watanabe et al., 2018). This delay may be caused by the loss of pre-assembled protein complexes at endocytic sites and slow recruitment of these proteins to the site of endocytosis. Consistent with this idea, recent studies indicate that ultrafast endocytosis is fast after a single stimulus but slows with high-frequency stimulation, consistent with a depletion of the depot of endocytic proteins (Delvendahl et al., 2016; Imig et al., 2020; Soykan et al., 2017). When the activity level increases dramatically, another mode of clathrin-independent endocytosis, bulk endocytosis, kicks-in to clear out excess membranes using the cytosolic Dyn1xB isoform (Clayton and Cousin, 2009) (Xue et al., 2011). Thus, we propose that clathrin-independent endocytosis is the predominant mechanism for synaptic vesicle retrieval, and the speed is determined by pre-assembly of endocytic proteins at synapses.

### Dynamin isoforms function in different pathways for synaptic vesicle regeneration

Dynamin 1 and 3 are highly expressed in brain and primarily involved in synaptic vesicle recycling (Ferguson et al., 2007; Raimondi et al., 2011). Furthermore, Dynamin 1 generates alternatively spliced isoforms, Dyn1xA and Dyn1xB, that only differ by short sequences at the C-termini. Our data indicate that Dyn1xA is required for ultrafast endocytosis, Dyn1xB functions during bulk endocytosis, and Dyn3 buds mature synaptic vesicles from the synaptic endosome.

Of the two abundant splice variants of Dyn1, only Dyn1xA can support ultrafast endocytosis. The two variants bind Syndapin 1 with similar affinity, yet Dyn1xB does not form puncta efficiently. This is not because Dyn1xB is phosphorylated, since more Dyn1xB is dephosphorylated at rest than Dyn1xA (Chan et al., 2010). Moreover, the phospho-deficient form of Dyn1xB S774/778A does not rescue the ultrafast endocytosis defect of *Dyn1,3* DKO neurons, suggesting that the phosphorylation status of this variant and binding to Syndapin 1 does not dictate whether it participates in ultrafast endocytosis.

The relevant difference between these two isoforms appears to be that Dyn1xA binds more tightly to Endophilin A. The capability to bind either Syndapin or Endophilin may be important for creating the multivalent interactions involved in phase separation. The proline-rich domain of Dynamin likely interacts with only one protein at a time (Anggono and Robinson, 2007), and therefore they will compete for binding in Dyn1xB. The C-terminal extension in Dy1xA introduces a second Endophilin binding site. Because Dynamin 1 exists in the cytosol as dimers and tetramers (Cocucci et al., 2014), Dyn1xA could act as the nucleating factor for multivalent interactions with itself and Endophilin A and Syndapin 1, and in turn Synaptojanin, and thereby increase network formation. Thus, the C-terminal extension of the xA variant would promote phase separation and clustering at the plasma membrane by binding Endophilin – interacting with both proteins is essential for Dyn1xA function.

Dyn1xB also plays an essential role in synaptic vesicle recycling, particularly when the activity level of synapses is high (Xue et al., 2011). Dyn1xB directly binds calcineurin (Xue et al., 2011). This interaction promotes dephosphorylation of Dyn1 in response to an increase in calcium at synaptic terminals and potentially switches the mode of endocytosis from ultrafast endocytosis to another clathrin-independent endocytosis, activity-dependent bulk endocytosis (Xue et al., 2011). Our results suggest that application of high potassium, which induces bulk endocytosis, increases the number of Dyn1xA puncta substantially, and each punctum contains more Dyn1xA, suggesting that docking of calcineurin on Dyn1xB allows cross-dephosphorylation of Dyn1xA and increases the amount of available Dyn1xA to mediate endocytosis. Thus, Dyn1xB likely plays a critical role in tuning endocytic capacity at synapses.

Dyn3 also contributes to synaptic vesicle endocytosis (Raimondi et al., 2011). In the *Dyn1* KO, endocytosis is severely impaired, but this phenotype can be reversed following a brief recovery period, suggesting other Dynamin isoforms may be able to mediate membrane fission in the absence of Dyn1. In fact, in the *Dyn1*,*3* DKO, the endocytic defect was more severe, indicating Dyn3 can function in the same pathway (Raimondi et al., 2011). However, *Dyn3* KO alone displayed no endocytic defect, and thus, it was unclear how Dyn3 supports synaptic vesicle endocytosis. Our data are consistent with previous experiments that endocytosis is normal in *Dyn3* KO and it is likely that Dyn3 only functions at the plasma membrane in the absence of Dyn1. However, synaptic vesicles are less efficiently generated from synaptic endosomes. In the wild type, synaptic endosomes are fully resolved into synaptic vesicles by 10 s after stimulation (Watanabe et al., 2014). In contrast, endosomes are still present at this time point in *Dyn3* KO neurons, and less than ∼50% of vesicles have been regenerated, suggesting that Dyn3 may be involved in the resolution of endosomes. However, further investigation is warranted to reveal the role of Dyn3 at endosomes.

### Syndapin 1 recruits Dyn1xA to the plasma membrane

Syndapin 1 is required for the recruitment of Dyn1xA to the plasma membrane. Two structural features make Syndapin 1 unique among the BAR protein family and are essential for dynamin localization at synapses. First, the SH3 domain of Syndapin 1 interacts with the Dyn1xA PRD at residues on either side of the phosphodomain (Figure S3B), making the affinity much higher than for a typical PRD – SH3 interaction (Anggono and Robinson, 2007). We found that this interaction is necessary for Dyn1xA localization to the plasma membrane. Second, the Syndapin 1 BAR domain is flexible and almost flattens out on membranes (Mahmood et al., 2019). This feature likely allows Syndapin 1 to interact with the flat plasma membrane without inducing curvature. In addition, the Syndapin BAR domain has a short amphipathic ‘wedge’ loop, which can be inserted into the lipid bilayer, possibly bending the membrane, to aid during pit formation (Rao et al., 2010; Wang et al., 2009). However, pit formation was normal when we introduced negatively charged residues into the wedge loop, even though Syndapin was no longer associated with the membrane. Thus, the BAR domain does not play a role in early steps of endocytosis but rather tethers Dyn1xA to the plasma membrane to accelerate neck cinching during ultrafast endocytosis.

### The kinetic control of endocytosis

How does stockpiling Dyn1xA accelerate endocytosis? The simplest model is that phase separation maintains the critical concentration of Dyn1xA at endocytic sites for efficient oligomerization and fission. *In vitro* experiments suggest that oligomerization (Stowell et al., 1999), phospholipid binding (Powell et al., 2000) and rate of membrane fission (Pucadyil and Schmid, 2008) are all enhanced when dynamin concentration is increased. This concentration is typically not achieved during clathrin-mediated endocytosis – dynamin gradually accumulates near endocytic pits and is recruited to the neck by other proteins such as Endophilin and Amphiphysin (Aguet et al., 2013; Cocucci et al., 2014; Macia et al., 2006; Merrifield et al., 2002; Taylor et al., 2011b, 2012). Phase separation bypasses such a recruitment phase, which normally takes seconds to tens of seconds (Cocucci et al., 2014; Taylor et al., 2012), and perhaps maintains Dyn1xA concentration at the critical concentration at all times near the endocytic sites for efficient oligomerization, phospholipid binding, and membrane fission.

In addition to enhancing Dyn1 activity, phase separation may allow Dyn1xA to participate in the early phase of endocytosis during cinching of the neck, not just in scission. Unlike Dyn2, which requires a narrow neck generated by multiple BAR domain proteins for it to function (Neumann and Schmid, 2013), Dyn1 can be recruited to shallow endocytic pits and promote the transition of shallow pits to deep invaginations (Liu et al., 2011). This process can bypass the requirement for BAR proteins in neck formation. Endocytosis occurs, albeit slowly, in the absence of the BAR-domain protein Endophilin A suggesting Dyn1 might be capable of acting without Endophilin (Watanabe et al., 2018). Thus, Dyn1 might assist and accelerate neck cinching. Indeed, when Dyn1 is recruited during the early phase of clathrin-mediated endocytosis, endocytosis is substantially faster than endocytic events relying solely on Dyn2 (Srinivasan et al., 2018). Thus, phase separation may recruit Dyn1xA to participate in an essential manner in neck formation with Synaptojanin and Endophilin A (Watanabe et al., 2018) during ultrafast endocytosis. Endocytic pits arrested on the plasma membrane in *Dyn1,3* DKO neurons have a wide opening at their base – similar to the pits observed in *Synaptojanin 1* KO and *Endophilin A* triple KO (Watanabe et al., 2018). By contrast, when the GTPase activity of dynamin, but not GTP-binding, is inhibited by the application of Dynasore (Macia et al., 2006) in wild-type neurons, the membrane gap at the base of endocytic pits is almost closed (Watanabe et al., 2013b). These data support *in vitro* experiments suggesting that the presence of dynamin is likely necessary for neck formation (Liu et al., 2011). Thus, phase separation can potentially accelerate ultrafast endocytosis by inducing rapid oligomerization, and membrane remodeling by Dyn1, Endophilin and Synaptojanin.

## Acknowledgements

We thank Pietro De Camilli for sharing mice and antibodies, Ira Milosevic for advice on antibodies, and Philip Robinson for discussion. We are also indebted to M. Delanoy and B. Smith at the Johns Hopkins Microscopy Facility for technical assistance in electron microscopy, Tyler Ogunmowo for the preparation of cells, and Grant F. Kusick for editing of the manuscript. S.W. and this work were supported by start-up funds from the Johns Hopkins University School of Medicine, Johns Hopkins Discovery funds, Johns Hopkins Catalyst award, the National Science Foundation (1727260), and the National Institutes of Health (1DP2 NS111133-01 and 1R01 NS105810-01A1) awarded to S.W, and German research council funded grants CRG958/A5, Exc257 and the Reinhard Koselleck project awarded to C.R.. S.W. is an Alfred P. Sloan fellow, a McKnight Foundation Scholar and a Klingenstein and Simons Foundation scholar. Y.I. was supported by JSPS. M.A.C. is supported by The Wellcome Trust (204954/Z/16/Z). E.M.J. is supported by the NIH grant NS034307, and is an Investigator of the Howard Hughes Medical Institute.

## Author Contributions

Y.I., S.R., and S.W. conceived the study and designed the experiments. C.R. and E.M.J. oversaw the pilot phase of the project. S.W. oversaw the overall research. Y.I., S.R., P.F., L.M.. F.Z., and B.S. performed the freezing experiments. Y.I., S.R., E.S., and S.W. collected and analyzed the data. S.N. and J.I. generated the scientific animation. E.B. and M.A.C. performed Western blots. Y.I., S.R., E.S., K.I., and T.T. generated DNA constructs for the study. Y.I and S.W. wrote the manuscript. All authors contributed to editing of the manuscript. C.R. and S.W. funded the research.

## STAR Methods

### EXPERIMENTAL MODEL AND SUBJECT DETAILS

All experiments were performed according to the rules and regulations of the National Institute of Health, USA, the UK Animal (Scientific Procedures) Act 1986, under Project and Personal Licence authority (Home Office project licence – 7008878), and Berlin, Germany authorities. Animal protocols were approved by committee of animal care, use of the Johns Hopkins University, the University of Edinburgh Animal Welfare and Ethical Review Body, and welfare committee of the Charite, Berlin. Primary cultures of mouse hippocampal neurons with the following genotypes were used in this study: C57/BL6-N; C57/BL6-J, *DNM3* KO (Raimondi et al., 2011), and *DNM1,3* DKO (Raimondi et al., 2011). The animals of the genotype *DNM1^+/+^*, *DNM3*^-/-^ and *DNM1^-/-^*, *DNM3*^-/-^ were generated by crossing *DNM1*^+/-^, *-3*^-/-^ animals. *DNM1^+/-^*, *Dnm3*^-/-^ animals were not included in the study to avoid any confound results due to the haploinsufficiency.

### Primary neuronal cultures

To prepare primary neuronal cultures, the following procedures were carried out. Newborn or embryonic day 18 (E18) mice of both genders were decapitated. The brain is dissected from these animals and placed on ice cold dissection medium (1 x HBSS, 1 mM sodium pyruvate, 10 mM HEPES, 30 mM glucose, and 1% penicillin-streptomycin).

For high pressure freezing, hippocampal neurons were cultured on a feeder layer of astrocytes. Astrocytes were harvested from cortices with treatment of trypsin (0.05%) for 20 min at 37 °C, followed by trituration and seeding on T-75 flasks containing DMEM supplemented with 10% FBS and 0.2% penicillin-streptomycin. After 2 weeks, astrocytes were plated onto 6-mm sapphire disks (Technotrade Inc) coated with poly-D-lysine (1 mg/ml), collagen (0.6 mg/ml) and 17 mM acetic acid at a density of 13 x 10^3^ cells/cm^2^. After 1 week, astrocytes were incubated with 5-Fluoro-2′-deoxyuridine (81 µM) and uridine (204 µM) for at least 2 hours to stop the cell growth, and then medium was switched to Neurobasal-A (Gibco) supplemented with 2 mM GlutaMax, 2% B27 and 0.2% penicillin-streptomycin prior to addition of hippocampal neurons. Hippocampi were dissected under a binocular microscope and digested with papain (0.5 mg/ml) and DNase (0.01%) in the dissection medium for 25 min at 37 °C. After trituration, neurons were seeded onto astrocyte feeder layers at density of 20 x 10^3^ cells/cm^2^. Cultures were incubated at 37 °C in humidified 5% CO_2_/95% air atmosphere. At DIV14-15, neurons were used for high pressure freezing experiments.

For fluorescence imaging, dissociated hippocampal neurons were seeded on 18-mm or 25-mm coverslips coated with poly-L-lysine (1 mg/ml) in 0.1 M Tris-HCl (pH8.5) at a density of 25-40 x 10^3^ cells/cm^2^. Neurons were cultured in Neurobasal media (Gibco) supplemented with 2 mM GlutaMax, 2% B27, 5% horse serum and 1% penicillin-streptomycin at 37 °C in 5% CO_2_. Next day, medium was switched to Neurobasal with 2 mM GlutaMax and 2% B27 (NM0), and neurons maintained thereafter in this medium. For biochemical experiments, dissociated cortical neurons were seeded on poly-L-lysine coated plates with Neurobasal media supplemented with 2 mM GlutaMax, 2% B27, 5% horse serum and 1% penicillin-streptomycin, at a density of 1 x 10^5^ cells/cm^2^. Next day, the medium was switched to Neurobasal medium with 2 mM GlutaMax and 2% B27, and neurons maintained in this medium thereafter. A half of the medium was refreshed every week.

### Expression constructs

All the plasmids used in this study and primers to make these construct was listed in Table S1. DNA cloning was performed using transformation into competent DH5*α* cells. For the dynamin rescue constructs, wild-type Dyn1xA, Dyn1xB and Dyn2 from human sequences were amplified from plasmids (generously provided by Pietro De Camilli’s lab) and fused by gibson assembly (NEW ENGLAND BioLabs) after a self-cleaving P2A site within a pFUGW derived plasmid, which encodes nuclear RFP controlled by the human synapsin 1 promoter (f(syn)NLS-RFP-P2A), to form f(syn)NLS-RFP-P2A-Dyn1xA/Dyn1xB/Dyn2. Using the same approach, we generated Dyn1xA S774/778A, S774/778D, Dyn1xB S774/778A.

For Syndapin 1 shRNA, oligos containing the target sequence were annealed with T4 DNA ligase (NEW ENGLAND BioLabs) and phosphorylated with T4 polynucleotide kinase (NEW ENGLAND BioLabs). The annealed oligos were ligated downstream of a human U6 promoter using BamHI and PacI restriction sites within the modified FUGW FUGW vector (f(U6).hSyn-NLS-RFP-WPRE) that further contained a human synapsin-1 promoter controlled NLS-RFP reporter construct (f(U6)Synapsin 1-shRNA.hSyn-NLS-RFP-WPRE). For the Syndapin 1 rescue constructs, mouse Syndapin 1 sequence was amplified and ligated into NheI and AscI restriction sites of f(syn)NLS-RFP-P2A (f(syn)NLS-RFP-P2A-Syndapin 1). f(syn)NLS-RFP-P2A-Syndapin 1 construct was linearized using primers with shRNA Syndapin 1 resistant sequence and re-circularized using In-Fusion HD cloning kit (TAKARA) (f(syn)NLS-RFP-P2A-Syndapin 1 shRNA resistant). For the Syndapin 1 I122D/M123E rescue construct, f(syn)NLS-RFP-P2A-Syndapin 1 shRNA resistant construct was amplified using primers with I122D/M123E mutations. The resulting PCR solution was digested with DpnI to degrade template DNA followed by ethanol precipitation. DNA was then transformed to competent DH5*α* cells. For the Syndapin 1 P434L rescue construct, f(syn)NLS-RFP-P2A-Syndapin 1 shRNA resistant construct was amplified using primers with P434L mutations and re-circularized using In-Fusion HD cloning kit (TAKARA).

To generate mCherry-Syndapin 1 with the shRNA resistant sequence (mCherry-shRNA resistant Syndapin 1), mouse Syndapin 1 sequence was amplified and combined with linearized mCherry construct using In-Fusion HD cloning kit. Then, mCherry-Syndapin 1 construct was linearized using primers with shRNA syndapin resistant sequence and re-circularized using In-Fusion HD cloning kit (TAKARA). To make mCherry-shRNA resistant Syndapin 1 122D/M123E and mCherry-shRNA resistant Syndapin 1 P434L, mCherry-shRNA resistant Syndapin construct was linearized using the primers with corresponding mutations and were re-circularized using In-Fusion HD cloning kit.

Dyn1xA-GFP was purchased from addgene. To generate Dyn1xA S774/778A-GFP and Dyn1xA-S774/778D GFP, Dyn1xA-GFP construct was linearized using the primers with corresponding mutations and were re-circularized using In-Fusion HD cloning kit. To generate Dyn1xA-GFP_mCherry-Syb2, rat Syb2 (addgene) and mCherry (addgene) sequence were amplified and combined with linearized Dyn1xA construct using In-Fusion HD cloning kit. As a result, mCherry-Syb2 sequence are inserted in to downstream of Dyn1xA-GFP sequence. To generate cytosolic tdTomato, tdTomato sequence was amplified from tdTomato vector (Clontech) and was ligated at the XhoI and KpnI digestion sites of pCAGGS construct by DNA ligation kit Solution I (TAKARA).

For pull-down assays, wild-type rat Dyn1xA-PRD (C-terminus residues 746-864), S774/778E and S774/778A mutants were generated by amplifying the required region from Dyn1aa-GFP (rat) in pEGFP-N1. The amplified product was inserted into pGEX4T-1 vector (Amersham Biosciences) using the restriction enzymes EcoRI and NotI (underlined in sequence). Subsequent modifications to generate S774/778E and S774/778A mutants used the QuickChange site-directed mutagenesis kit (Stratagene) and were confirmed by DNA sequencing (Anggono et al., 2006). Wild-type rat Dynamin-1xb C-terminus (residues 746-851) was generated by amplifying the required region from Dyn1ab in a pCR3.1 expression vector (Cao et al 1998). Similar to Dyn1xA, the amplified product was inserted into a pGEX-4T-1 using the restriction enzymes EcoRI and NotI (Xue et al., 2011). To generate Dyn1xA-PRD S851/857A and Dyn1xA-PRD S851/857E, Dyn1xA-PRD was linearized using the primers with corresponding mutations and were re-circularized using T4-ligase (Promega, M1801). Dyn1xB-PRD S774/778A and Dyn1xB-PRD S774/778E were generated via the digestion of Dyn1xA-PRD S774/778A or Dyn1xB-PRD S774/778E with BamHI. This excised a sequence encoding the mutated residues (cut sites in multi-cloning site and between G806/S807). This fragment was ligated into wild-type Dyn1xB-PRD previously digested with BamHI. An identical protocol as followed for the Dyn1xA-PRD S774/778/851/857A and Dyn1xA-PRD S774/778/851/857E mutants, except the corresponding fragment was ligated into BamHI-digested Dyn1xA-PRD S851/857A or Dyn1xA-PRD S851/857E respectively. The correct insertion of the fragment was confirmed by Sanger sequencing, which was also the case for site-directed mutagenesis.

For GFP knock-in constructs, oligos containing the 20-bp target sequences were annealed at gradient temperature from 95 °C to 20 °C for 30 min. The annealed oligos were ligated into the BbsI sites of pORANGE vector (addgene) by DNA ligation kit Solution I (TAKARA). Donor sequence, consisting of GFP and two Cas9 target sites flanking GFP, was PCR amplified from pEGFP-N1 plasmid and cloned into the HindIII and XhoI site of pORANGE vector containing the target sequence. First, mCherry was PCR amplified from pmCherry-N1 plasmid (Clontech) with mC-F and mC-R primers and ligated into the BamHI and EcoRI site of pFUGW-spCas9 replacing Cas9, generating pFUGW-mCherry. Then, knock-in cassettes containing the U6 promoter, gRNA, and donor DNA was PCR amplified from pORANGE constructs and cloned into PacI site of pFUGW vector using In-Fusion HD cloning kit.

### Lentivirus production and infection

For lentivirus production, HEK293T cells were maintained in DMEM supplemented with 10% FBS and 0.2% penicillin-streptomycin. One day after plating 6.5 × 10^!^ cells in T75 flask (Corning), medium was replaced with Neurobasal-A media supplemented with 2 mM GlutaMax, 2% B27 and 0.2% penicillin-streptomycin. Cells were transfected using polyethylenimine with a pFUGW plasmid encoding the RFP co-infection marker and insert of interest, and two helper plasmids (pHR-CMV8.2 deltaR and pCMV-VSVG) at a 4:3:2 molar ratio. Three days after transfection, supernatant was collected and concentrated to 20-fold using Amicon Ultra-15 10K centrifuge filter (Millipore). ChetaTC and dynamin rescue viruses were added to each well of neurons at DIV 3 with 15 µl per well (12-well plates). Syndapin 1 shRNA virus was added to each well of neurons at DIV 3 with 5 µl per well (12-well plates, 75 K neurons / well). Syndapin 1 rescue viruses were added to each well of neurons at DIV 3 with 5 µl per well (12-well plates, 75K neurons / well) or 20 µl per well (6-well plates, 300 K neurons / well). In all cases, infection efficiency of over 95% was achieved. Neurons were fixed or used for high pressure freezing at DIV13-16.

### Transient transfection of neurons

For transient protein expression, neurons were transfected at DIV13-14 using Lipofectamine 2000 (Invitrogen) in accordance with manufacture’s manual. Before transfections, a half of medium in each well was transferred to 15 mL tubes and mixed with the same volume of fresh NM0 and warmed to 37 °C in CO_2_ incubator. This solution later served as a conditioned medium. Briefly, 0.25-2 µg of plasmids was mixed with 2 µl Lipofectamine in 100 µl Neurobasal media and incubated for 20 min. For Dyn1 expression, 0.5 µg of constructs were used to reduce the expression level. For tdTomato expression, 2.0 µg of constructs were used. For Syndapin 1 expression, 0.25 µg of constructs were used in the Syndapin 1 knock-down neurons. The plasmid mixture was added to each well with 1 ml of fresh Neurobasal media supplemented with 2 mM GlutaMax and 2% B27. After 4 hours, medium was replaced with the pre-warmed conditioned media. Neurons were incubated for less than 20 hours and fixed for imaging or subjected to live imaging. For GFP knock-in experiments, hippocampal neurons were transfected at DIV 3 with pORANGE construct (1.5 µg) and pmCherry-N1 (0.5 µg) as cell-fill using Lipofectamine 2000 and fixed at DIV15-18.

### Sequencing of genomic sites of GFP integration

Genomic DNA was isolated from the neurons at DIV15-17 using DNeasy Blood & Tissue Kit (Qiagen). Genomic PCR was performed to amplify the 5’ and 3’ junctions of the integrated GFP by KOD Hot Start DNA Polymerase (Novagen) using the primers listed in Table S1. PCR products were separated by agarose gel agarose gel electrophoresis and the fragments were purified using QIAquick Gel extraction kit (Qiagen). Purified DNA was cloned into pCR4Blunt TOPO vector using Zero Blunt TOPO PCR Cloning Kit (Invitrogen). The sequences of 18 clones were analyzed.

### Immunofluorescence staining

For immunofluorescence, 125 k neurons were seeded on 18 mm poly-L-lysins (1 mg/mL, overnight) coated coverslips (Thickness 0.09-0.12 mm, Caroline Biological) in 12-well plate (Corning). Neurons were fixed at DIV14-15 with pre-warmed (37 °C) 4% paraformaldehyde and 4% sucrose in PBS for 20 min and permeabilized with 0.2% Triton X-100 in PBS for 8 min at room temperature. After blocking with 1% BSA in PBS for 30 min, cells were incubated with primary antibodies diluted in 1% BSA/PBS overnight at 4 °C, followed by appropriate secondary antibodies diluted 1:400 in 1% BSA/PBS for 1 hour at room temperature. Samples were mounted in ProLong Gold Antifade Mountant (Invitrogen) and stored at 4 °C until imaging.

### Potassium stimulation

Neurons (DIV 15) were stimulated using 90 mM KCl as described previously (Wu et al., 2014). After a rinse with Tyrode buffer without Ca^2+^ (119 mM NaCl, 2.5 mM KCl, 2 mM MgCl_2_, 25 mM Hepes, and 30 mM glucose, pH 7.4) prewarmed to 37 °C, the same solution containing 2 mM Ca^2+^ and either 2.5 mM or 90 mM K^+^ was applied to neurons (37 °C) for 90 s. The ionic balance was maintained by reducing the Na^+^. Neurons were fixed immediately by application of pre-warmed (37 °C) 4% paraformaldehyde and 4% sucrose in PBS for 20 min.

### Confocal microscopy imaging and analysis

For fluorescence imaging, all samples were imaged using a confocal microscope Zeiss LSM880 (Carl Zeiss). Fluorescence was acquired using a 63x objective lens (NA = 0.55) at 2048×2048 pixel resolution with the following settings: pixel Dwell 1.02 μs and pin hole size at 2 airy unit. For experiments comparing fluorescence intensity, these settings were left constant between samples. Snapshots were captured at the cell body first and then 3-7 different locations along the axon of the same neurons, typically covering 1-3 mm length from each neuron. All Dyn1-GFP transfected neurons in each coverglass were imaged. Axons were distinguished from dendritic processes based on their morphology (thin and lacking spines). If GFP signals were aberrantly saturated throughout the processes, they were dismissed to avoid overexpression artifact. Apparently dying neurons and glia cells were also excluded. We quantified all presynaptic varicosities along axons in each image. Presynaptic regions were confirmed with the Syb2-Alexa647 signals. The Syb2-Alexa647 signals were used to define the boundaries of regions-of-interest (ROIs) for quantifications, and all Dyn1-GFP and mCherry-Syndapin 1 signals within ROIs were measured as the total signals at each synapse. To define puncta within the boutons, we applied Gaussian smoothing (*σ* = 0.2) on images, performed threshold cut-off on the average fluorescence intensity of Dyn1xA and Syndapin 1 in the intersynaptic axonal regions. The circumference of each punctum adjacent to or within Syb2 signals were delineated and set as ROIs, and fluorescence intensity measured. For the 90 mM KCl stimulation experiments, the diameter of puncta appeared larger at 90 s after the stimulation, and thus, integrated raw intensity within the ROI of the puncta was measured. The background signals were subtracted using the same size of ROI but 40-60 pixels away from the original ROI and outside of neural processes. For the quantification of the colocalization between Dyn1xA and Syndapin 1 puncta, ROIs of overlapped Dyn1xA and Syndapin 1 were counted. Fluorescence intensity was normalized to the average signal within the corresponding cell body of each axon. False colored images were made by applying LUT colors and ratio signal intensities are measured by calibrate bar function in Fiji.

### Live imaging and analysis

For live-cell imaging, 250 k neurons were seeded on 25 mm coverslips (thickness 0.13-0.17 mm, Carolina Biological) in 6-well plate (Corning). Dyn1xA-GFP and mCherry-Syb2 were expressed on the same construct using pDyn1xA-GFP_P2A_mCherry-Syb2 construct by transient transfection. For the digitonin treatment, 10 mg/mL of digitonin (Sigma-Aldrich) was prepared in MilliQ water and diluted into final concentration 5 μg/mL or 500 μg/mL in NM0. For the 1,6-Hexanediol treatment, 4 % of 1,6-Hexanediol was directly dissolved in NM0. For imaging, cover slips were transferred to live cell round chamber (ALA science) filled with NM0. FK506 (Tocris) or CHIR99021 (Sigma-Aldrich) were dissolved in DMSO and added to the imaging chamber to achieve the final concentrations of 2 µM and 10 µM, respectively, 30 min before imaging. 0.05 % of DMSO were added to the chamber for control experiments. All live imaging experiments were carried out on a Zeiss LSM880 confocal microscope at 37 °C in humidified 5% CO_2_/95% air atmosphere, and live imaging data was acquired by raster scan.

For photobleaching experiments, 33 frames were collected with a time interval of 2.5 s. After 3 frames, the maximum laser power was applied to bleach fluorescence. The 4th frame was started immediately after bleaching. 30 additional frames were collected after bleaching. A circular ROI was placed around each Dyn1xA punctum found adjacent to the mCherry-Syb2 signal, and Dyn1xA-GFP signals within the ROI was photobleached with a 488-nm laser. For quantifications, average fluorescence intensity was measured from each ROI and a random region outside of the cell, which was used for the background subtraction. A circular ROI of the same size was used to measure signals from an unbleached region to measure the degree of imaging-related photobleaching. The fluorescence intensity at the ROIs was normalized to the intensity before bleaching (3rd frame) and after bleaching (4th frame). To correct signal loss due to the bleaching during the imaging, the normalized fluorescence recovery over time, *F*(*t*), was then fitted to an exponential function:

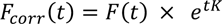

where the corrected normalized fluorescence over time is *F*_corr_(*t*), time after the photo bleaching is *t*, and *K* is the rate constant acquired from the unbleached region and measured exponential curve fitting tool in Fiji.

The *F*_corr_(*t*), was then fitted to an exponential function using GraphPad Prism (v8);

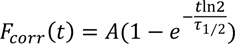

where the recovery is *A*, half-recovery time is *τ*_*τ*1⁄2_..

The diffusion coefficient, D, was fitted to an,

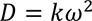

where the rate constant acquired from the bleached region is *k* and *ω* is the diameter of the bleached region.

For the cytosolic extraction experiments with digitonin, a total of 17 frames were collected with a time interval of 5 s: 5 frames were collected prior to the application of digitonin (500 μg/mL in NM0)(Sigma-Aldrich), and additional 12 frames were collected thereafter. For the 1,6,-hexanediol treatment, a total of 12 frames were collected with a time interval of 30 s: 2 frames before the addition of 5 μg/mL of digitonin and 4 % 1,6,-hexanediol (Sigma-Aldrich) in NM0 and 10 frames afterwards. For the purpose of imaging-related photobleaching corrections, the same number of frames were collected at additional regions distant from the imaged before the experiments. For quantifications, the average fluorescence intensities before and after the treatment were measured within the same ROIs, which were selected as described in the previous section: one ROI covering the entire Syb2 signals and another ROI along the circumference of the Dyn1xA-GFP puncta. For presentations, the brightness and the contrast in each image were adjusted in Fiji and cropped in Adobe photoshop 2021 (Adobe).

### Protein expression and GST-pulldown assay

Isolated nerve terminals, or synaptosomes, were prepared from rat brains of both sexes (Cousin and Robinson 2000). GST fusion proteins were expressed in bacteria at 19 °C for 16-18 hours in 500 ml cultures of supermedia (5g/L NaCl, 25g/L yeast extract (Alfa Aesar J60287), 15g/L Tryptone (Merk 1.11931), pH7.4) and coupled to glutathione-Sepharose beads (GE healthcare, 17-0756-01).

Nerve terminals were solubilized for 5 minutes at 4 °C in 25 mM Tris, pH 7.4, with 1% Triton X-100, 150 mM NaCl, 1 mM EGTA, 1 mM EDTA, 1 mM PMSF and protease inhibitor cocktail (Sigma, P58849) and centrifuged at 20,442 g for 5 minutes at 4 °C. The subsequent supernatant was incubated with GST-fusion proteins for 2 hours at 4 °C under constant rotation. After washing in lysis buffer (including a 500 mM NaCl wash), beads were washed twice in cold 20 mM Tris (pH 7.4) and boiled in SDS sample buffer (67 mM SDS, 2 mM EGTA, 9.3% glycerol, 12% β-mercaptoethanol, 0.7 mg/mL Bromophenol blue, 67 mM Tris-HCl, pH 6.8). The released proteins were separated by SDS-PAGE and analyzed by Western blotting. Primary antibodies were incubated with the blots overnight at 4°C under constant rotation (anti-endophillin, sc10874, 1:3000; anti-syndapin/PACSIN1, ab137390, 1:5000; anti-endophilin A2, a gift from Pietro De Camilli (Milosevic et al., 2011). IRDye secondary antibodies (800CW anti-goat IgG, #926-32214 and 680RD anti-rabbit, #926-68073 both 1:10000) were applied in Odyssey blocking PBS buffer for 2 hours at room temperature (LI-COR Biosciences, Nebraska, USA). Blots were visualized using a LiCOR Odyssey fluorescent imaging system, with band densities quantified using Fiji (ImageJ) software. Both syndapin and endophilin bands were normalized to the GST fusion protein band revealed by Ponceau-S staining.

### Evaluations of Syndapin 1 shRNA efficiency

To test the efficiency of Syndapin 1 KD and the amount of overexpression, cultured hippocampal neurons were harvested, lysed by addition of lysis buffer (50 mM Tris pH 8.0 and 1% SDS containing cOmplete Mini Protease Inhibitor (Roche)) and boiling at 95°C for 5 min. Lysates were centrifuged at 15,000 x g for 15 min at 4 °C and the supernatants were collected. Proteins were separated by SDS– PAGE and transferred onto Immobilon-FL membranes (Millipore). After blocking with 5% skim milk in PBS containing 0.05% Tween-20 (PBST) for 30 min, membranes were incubated with anti PACSIN 1 rabbit polyclonal antibody diluted at 1:1000 in 3% BSA/PBST overnight at 4°C, followed by IRDye secondary antibodies (LiCor) diluted in 1:10,000 in 3% BSA/PBST for 45 min at room temperature. Visualization was performed using LiCor Odyssey Clx and quantification was done by Image Studio from LiCor.

### Flash-and-freeze experiments

For flash-and-freeze experiments (Watanabe et al., 2013a, 2014). sapphire disks with cultured neurons (DIV14-15) were mounted in the freezing chamber of the high-pressure freezer (HPM100 or EM ICE, Leica), which was set at 37 °C. The physiological saline solution contained 140 mM NaCl, 2.4 mM KCl, 10 mM HEPES, 10 mM Glucose (pH adjusted to 7.3 with NaOH, 300 mOsm, 4 mM CaCl_2_, and 1 mM MgCl_2_. Additionally, NBQX (3 mM) and Bicuculline (30 mM) were added to suppress recurrent network activity following optogenetic stimulation of neurons. To minimize the exposure to room temperature, solutions were kept at 37 °C water bath prior to use. The table attached to the high-pressure freezer was heated to 37 °C while mounting specimens on the high-pressure freezer. The transparent polycarbonate sample cartridges were also warmed to 37 °C. Immediately after the sapphire disk was mounted on the sample holder, recording solution kept at 37 °C was applied to the specimen and the cartridge was inserted into the freezing chamber. The specimens were left in the chamber for 30 s to recover from the exposure to ambient light. We applied a single light pulse (10 ms) to the specimens (20 mW/mm^2^). This stimulation protocol was chosen based on the results from previous experiments showing approximately 90% of cells fire at least one action potential (Watanabe et al., 2013b). The non-stimulation controls for each experiment were always frozen on the same day from the same culture. We set the device such that the samples were frozen at 1 or 10 s after the initiation of the first stimulus. For ferritin-loading experiments, cationized ferritin (Sigma-Aldrich) was added in the saline solution at 0.25 mg/ml. The calcium concentration was reduced to 1mM to suppress spontaneous activity during the loading. The cells were incubated in the solution for 5 min at 37 °C. After ferritin incubation, the cells were immersed in the saline solution containing 4 mM Ca^2+^. For dynamin experiments, all samples were incubated with 1 µM TTX for overnight to block spontaneous network activity and reduce the number of pits arrested on the plasma membrane prior to flash-and-freeze experiments (Raimondi et al., 2011; Wu et al., 2014).

Following high-pressure freezing, samples were transferred into vials containing 1% osmium tetroxide (EMS),1% glutaraldehyde (EMS), and 1% milliQ water, in anhydrous acetone (EMS). The freeze-substitution was performed in an automated freeze-substitution device (AFS2, Leica) with the following program: −90C for 5–7 h, 5C per hour to −20 °C, 12 hours at −20 °C, and 10°C per hour to 20 °C. Following *en bloc* staining with 0.1% uranyl acetate and infiltration with plastic (30% for 3 hours, 70% for 4 hours, and 90% for overnight), the samples were embedded into 100% Epon-Araldite resin (Araldite 4.4 g, Epon 6.2 g, DDSA 12.2 g, and BDMA 0.8 ml) and cured for 48 hours in a 60 °C oven. Serial 40-nm sections were cut using a microtome (Leica) and collected onto pioloform-coated single-slot grids. Sections were stained with 2.5% uranyl acetate before imaging.

### Electron microscopy

Ultrathin sections of samples were imaged at 80 kV on the Philips CM120 at 93,000x magnification or on the Hitachi 7600 at 150,000x. At these magnifications, one synapse essentially occupies the whole screen, and thus, with the bidirectional raster scanning of a section, it is difficult to pick certain synapses, reducing bias while collecting the data. In all cases, microscopists were additionally blinded to specimens and conditions of experiments. Both microscopes were equipped with an AMT XR80 camera, which is operated by AMT capture software v6. About 120 images per sample were acquired. If synapses do not contain a prominent postsynaptic density, they were excluded from the analysis – typically a few images from each sample fall into this category.

### Analysis of electron micrographs

Electron micrographs were analyzed using SynpasEM. Briefly, images were pooled into a single folder from one set of experiments, randomized, and annotated blind. Using custom macros (Watanabe et al., 2020), the x-and y-coordinates of the following features were recorded in Fiji and exported as text files: plasma membrane, postsynaptic density, synaptic vesicles, large vesicles, endosomes, and pits. By visual inspection, large vesicles were defined as any vesicle with a diameter of 60-100 nm. Endosomes were distinguished by any circular structures larger than large vesicles or irregular membrane-bound structures that were not mitochondria or endoplamic reticulum. Late endosomes and multivesicular bodies were not annotated in this study. Pits were defined as smooth membrane invaginations within or next to the active zone, which were not mirrored by the postsynaptic membranes. After annotations, the text files were imported into Matlab (MathWorks). The number and locations of vesicles, pits, endosomes were calculated using custom scripts (Watanabe et al., 2020). The example micrographs shown were adjusted in brightness and contrast to different degrees, rotated and cropped in Adobe Photoshop (v21.2.1) or Illustrator (v24.2.3). Raw images and additional examples are provided in Figshare.com/projects/dynamin_phase_separation. The macros and Matlab scripts are available at https://github.com/shigekiwatanabe/SynapsEM.

### Statistical analysis

Detailed statistical information was summarized in table S2. All EM data are pooled from multiple experiments after examined on a per-experiment basis (with all freezing on the same day); none of the pooled data show significant deviation from each replicate. Sample sizes were based on our prior flash-and-freeze experiments (∼2-3 independent cultures, over 200 images), not power analysis. An alpha was set at 0.05 for statistical hypothesis testing. Normality was determined by D’Agostino-Pearson omnibus test. Comparisons between two groups were performed using a two-tailed Welch two-sample t-test if parametric or Wilcoxon rank-sum test and Mann-Whiteney Test if nonparametric. For groups, full pairwise comparisons were performed using one-way analysis of variance (ANOVA) followed by Holm-Šídák’s multiple comparisons test if parametric or Kruskal-Wallis test followed by Dunn’s multiple comparisons test if nonparametric. All statistical analyses were performed and all graphs created in Graphpad Prism (v8).

All fluorescence microscopy data were first examined on a per-experiment basis. For Figure 1B, 1D, 1F, 1H, 1J-N, 2C, 2D, 2F, 2N, 2L, 3B-D, 3G-I, 3H, 3L, 4C, 5B, 5D, 5F, 5H, 5J-N, 6C, 6D, 6G, 6J-M, 7B, 7D, 7F, 7H, 7J, 7K-N, S1A, S1C, S2B, S2D, S2F, S2H-2L, S5A-C, S6B and S7C-E, the data were pooled; none of the pooled data show significant deviation from each replicate. Sample sizes were ∼2-3 independent cultures, at least 50 synapses from 4 different neurons in each condition. An alpha was set at 0.05 for statistical hypothesis testing. The skewness was determined by Pearson’s skewness test in GraphPad Prism (v8). Since data were all nonparametric, Mann–Whitney test or Kruskal–Wallis test were used. p-values in multiple comparison were adjusted with Bonferroni correction. Confidence levels were shown in each graph. All statistical analyses were performed, and all graphs created in Graphpad Prism (v8).

### Scientific Animation

The axon and dendrite were modeled using 3DEM images from previous studies as reference (Bromer et al., 2018; Harris et al., 2015). The bouton and its contents were modeled by referring to the 3D model of synaptic architecture (Wilhelm et al., 2014). The structure and composition of a synaptic vesicle were based on the molecular model generated by previous studies (Takamori et al., 2006; Tao et al., 2018). Structures of proteins in the animation were obtained from the following files from the Protein Data Bank −2KOG for synaptobrevin (VAMP-2), 2R83 for synaptotagmin-1, 6BAT for glutamine transporter, 5VOX for V-ATPase, 1PK8 for synapsin I, 3RAB for RAB3A, 3C98 for syntaxin-1A, 6MDM for SNAP-25, 5GJV for voltage-gated calcium channel, 5W5C for complexin-1, 3C98 for munc18a, 5UE8 for munc13a, 4PE5 for NMDA receptor, 3KG2 for AMPA receptor, 2X3X for syndapin-1, 1ZWW and 3IQL for endophilin-A1, 1I9Z and 3LWT for synaptojanin-1, 6DLU, 6DLV and 5A3F for dynamin-1 and 3IYV for clathrin. The 3D structure file of glutamine was acquired from PubChem (CID 5961). Calcium and protons were represented as spheres. The axon, dendrite, bouton, the membranes, and the missing regions of proteins were modeled in the 3D graphics application, Autodesk Maya (https://www.autodesk.com/products/maya/). Surface meshes of protein structures were obtained from the molecular visualization application, UCSF Chimera (Pettersen et al., 2004), and imported into Autodesk Maya for animation. The overall ultrafast synaptic vesicle cycling process depicted in this animation was built on information from previous studies (Chanaday et al., 2019; Watanabe et al., 2013b). The dynamic movement of synaptic vesicles at the active zone was inferred from a previous study (Neher and Brose, 2018). The details of calcium triggered fusion mediated by SNAREs, including formation of a primed complex with synaptotagmin and complexin facilitated by Munc proteins, were derived from previous studies (Bai et al., 2016; Chang et al., 2018; Palfreyman and Jorgensen, 2017; Wang et al., 2019; Zhou et al., 2017). Research that was referred to for endocytic pit formation, membrane bending by syndapin and endophilin, synaptojanin hydrolysis and membrane fission by dynamin, are published in previous studies (Chang-Ileto et al., 2011; Chappie et al., 2011; Colom et al., 2017; Frost et al., 2008; Mahmood et al., 2019; Mim et al., 2012; Watanabe et al., 2018). Clathrin-mediated endosomal budding was referenced from previous studies (Morris et al., 2019; Paraan et al., 2020), and vATPase inactivity in the presence of a clathrin coat was based on findings from a previous study (Farsi et al., 2018). The animated scenes were rendered into a series of images and imported into the post production application, Adobe After Effects (https://www.adobe.com/products/aftereffects) for compositing, and including annotation and audio to create the final video.

### Source data availability

Source data are provided with the manuscript and through https://data.mendeley.com/datasets/bkn78y232y/draft?preview=1. All raw images are available upon request.

**Supplemental Information:**

## STAR METHODS

### KEY RESOURCES TABLE

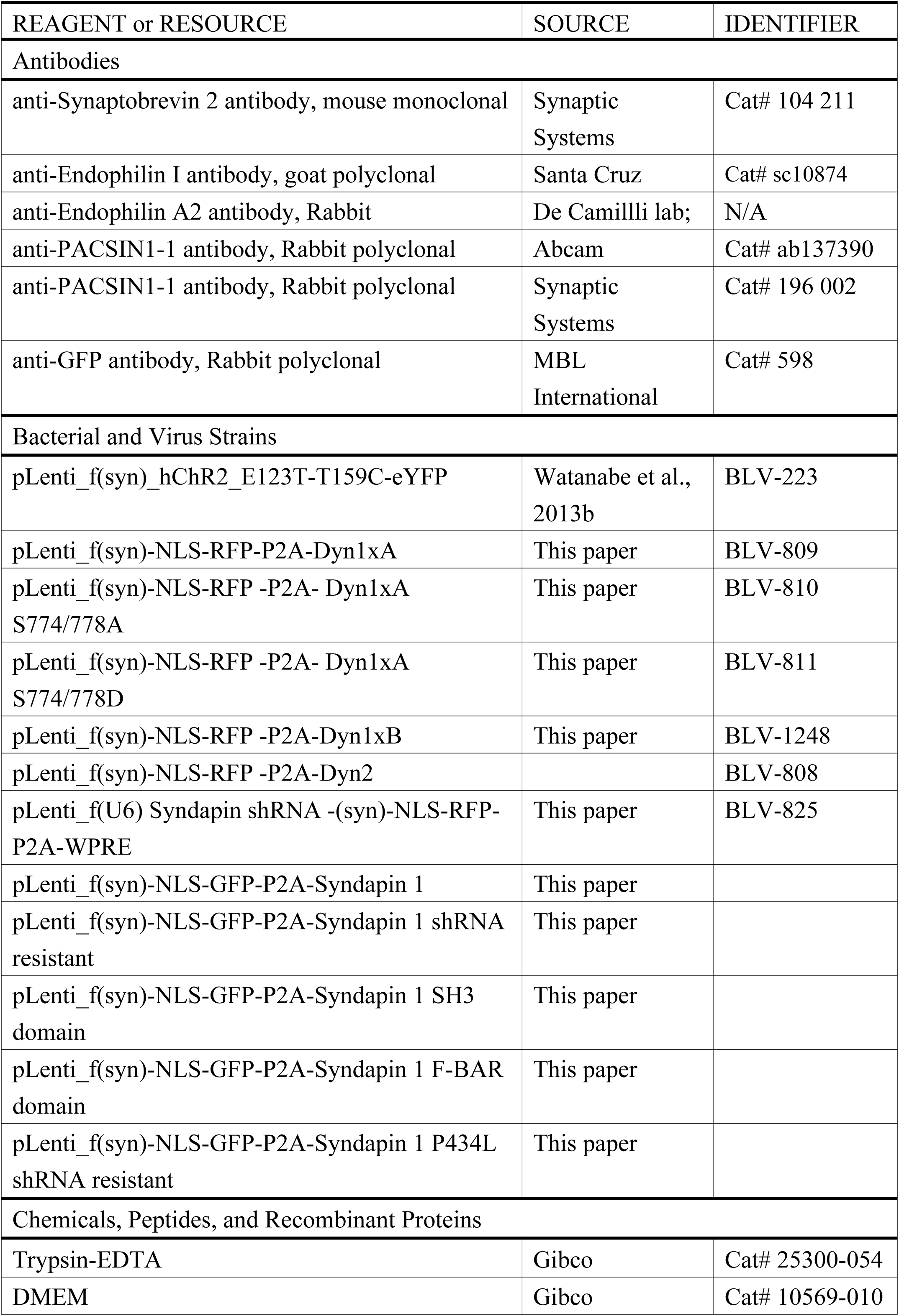
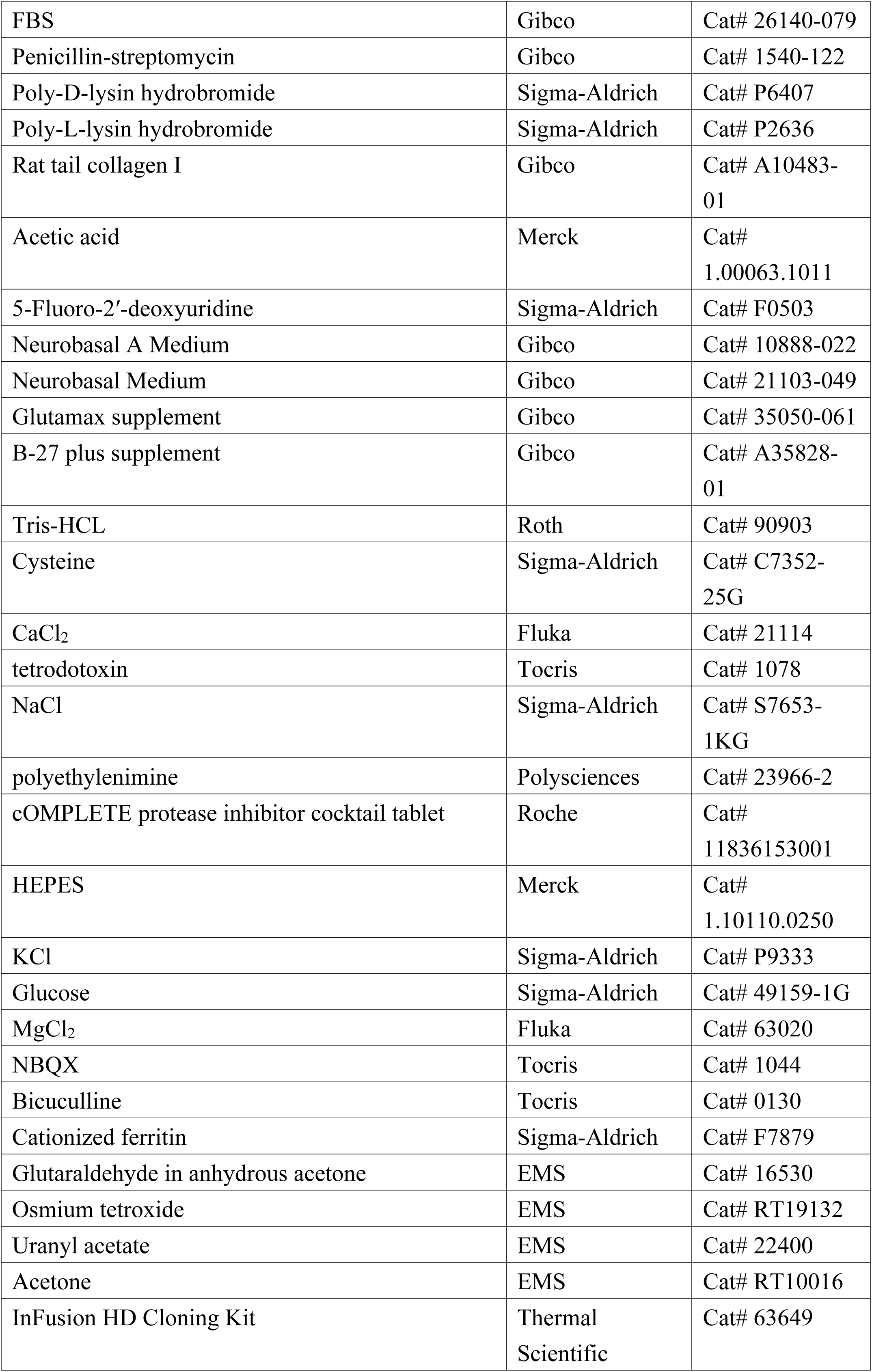
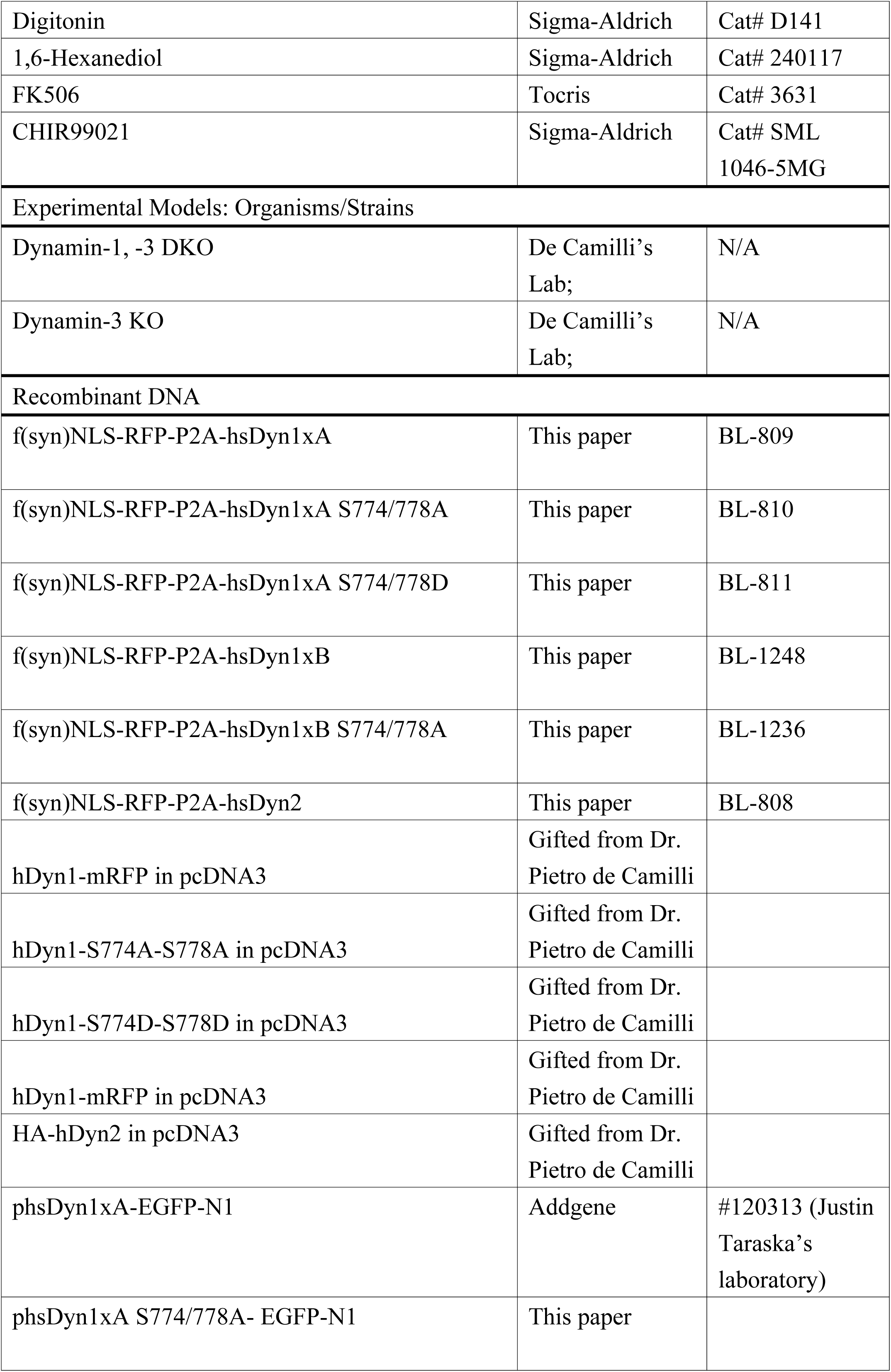
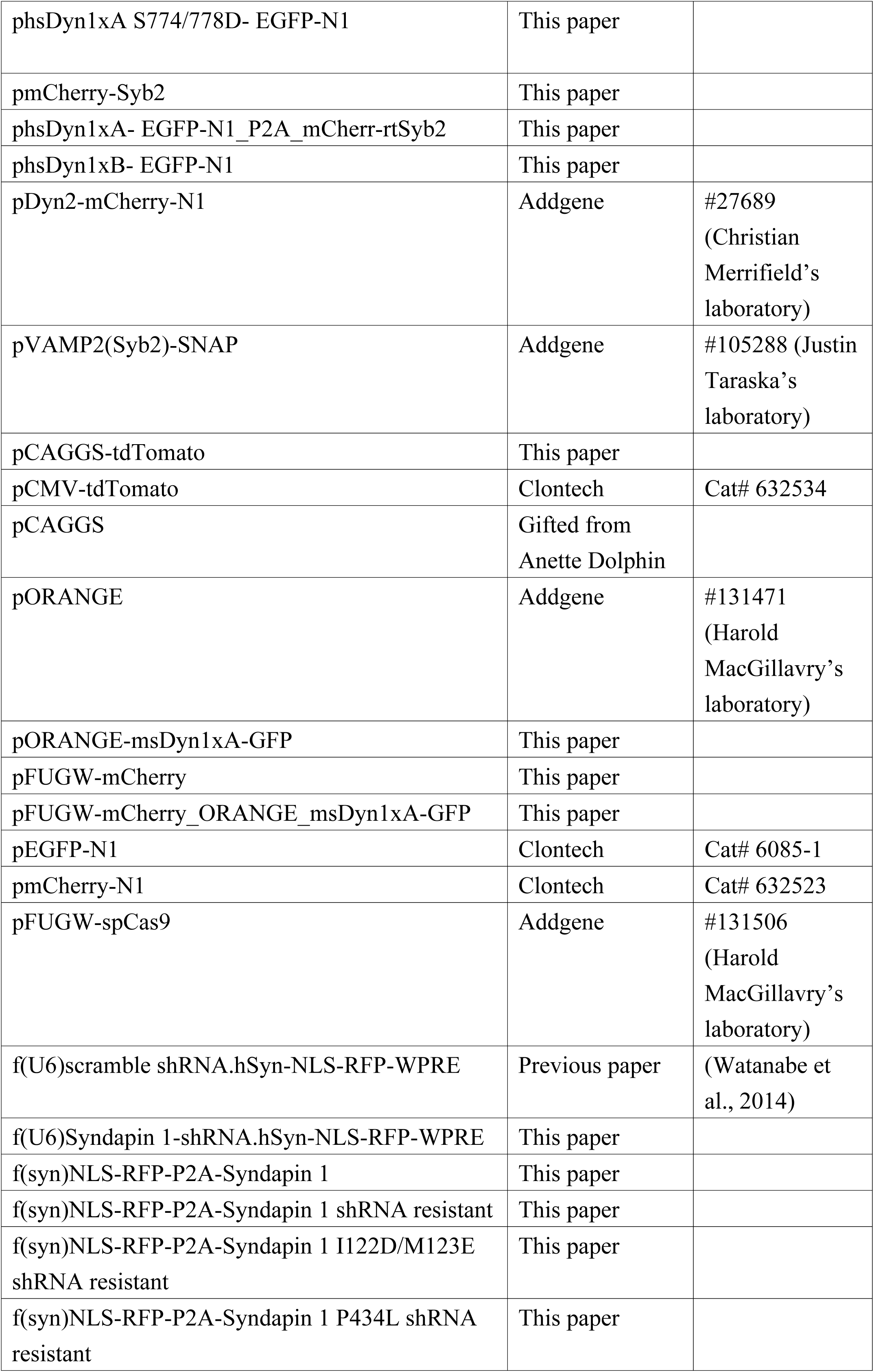
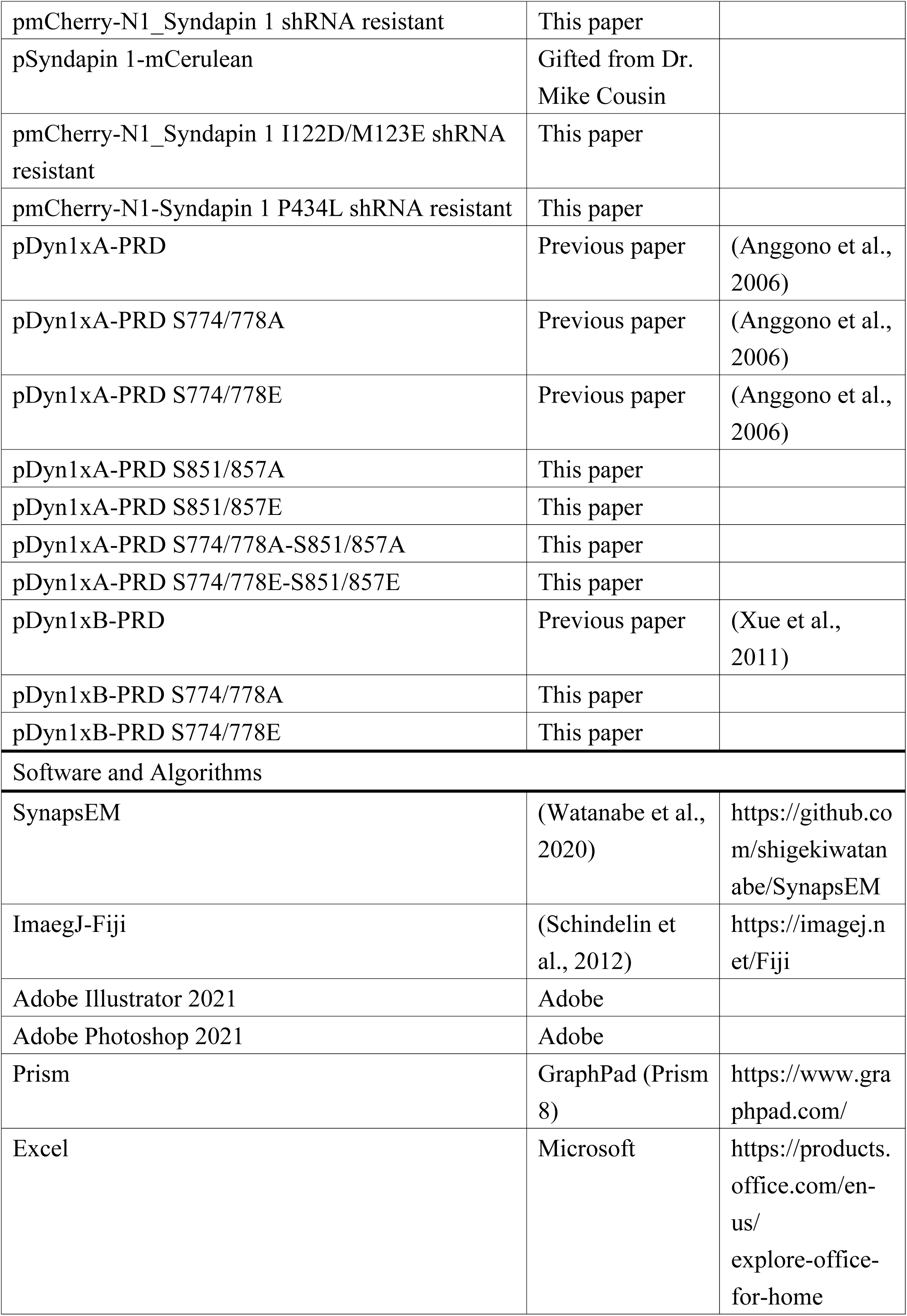
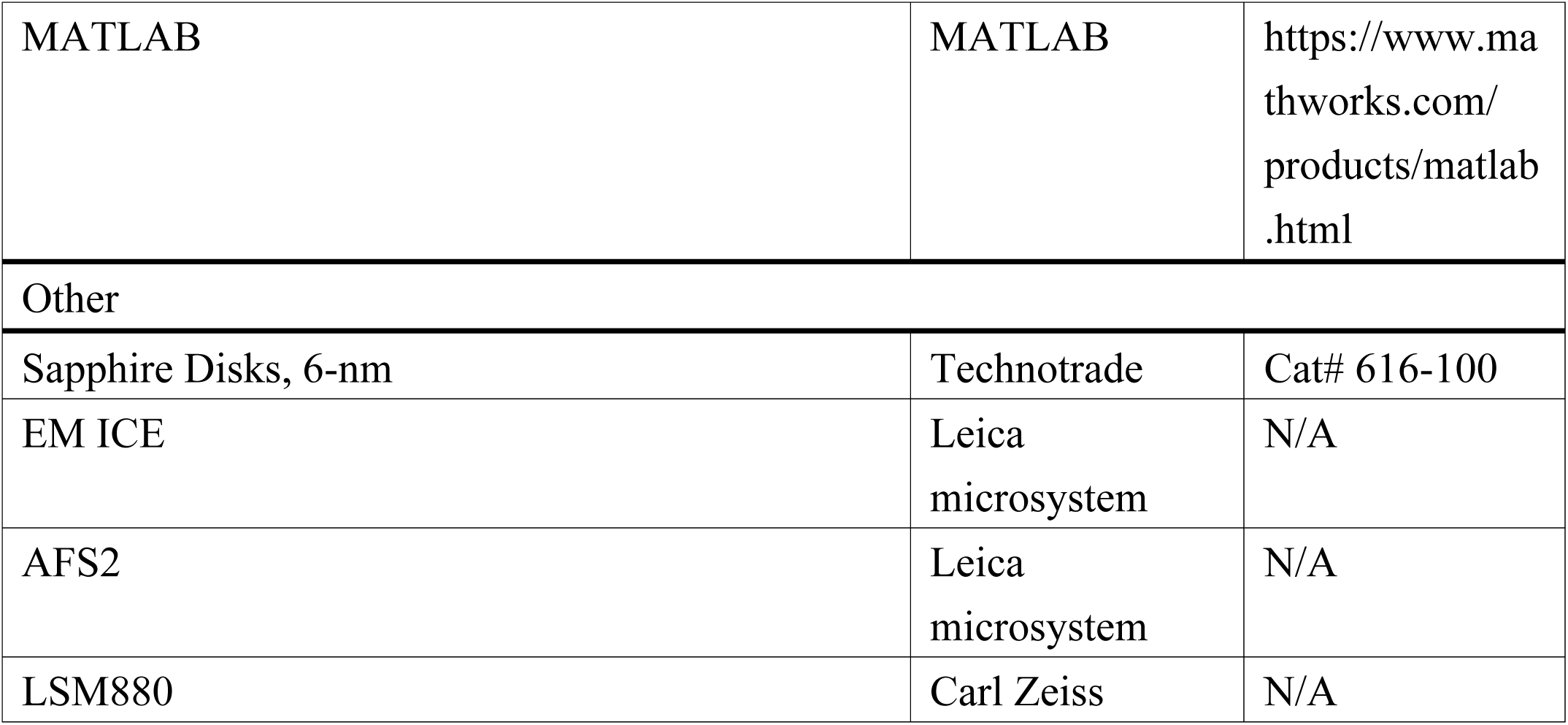

---

**Figure S1.**
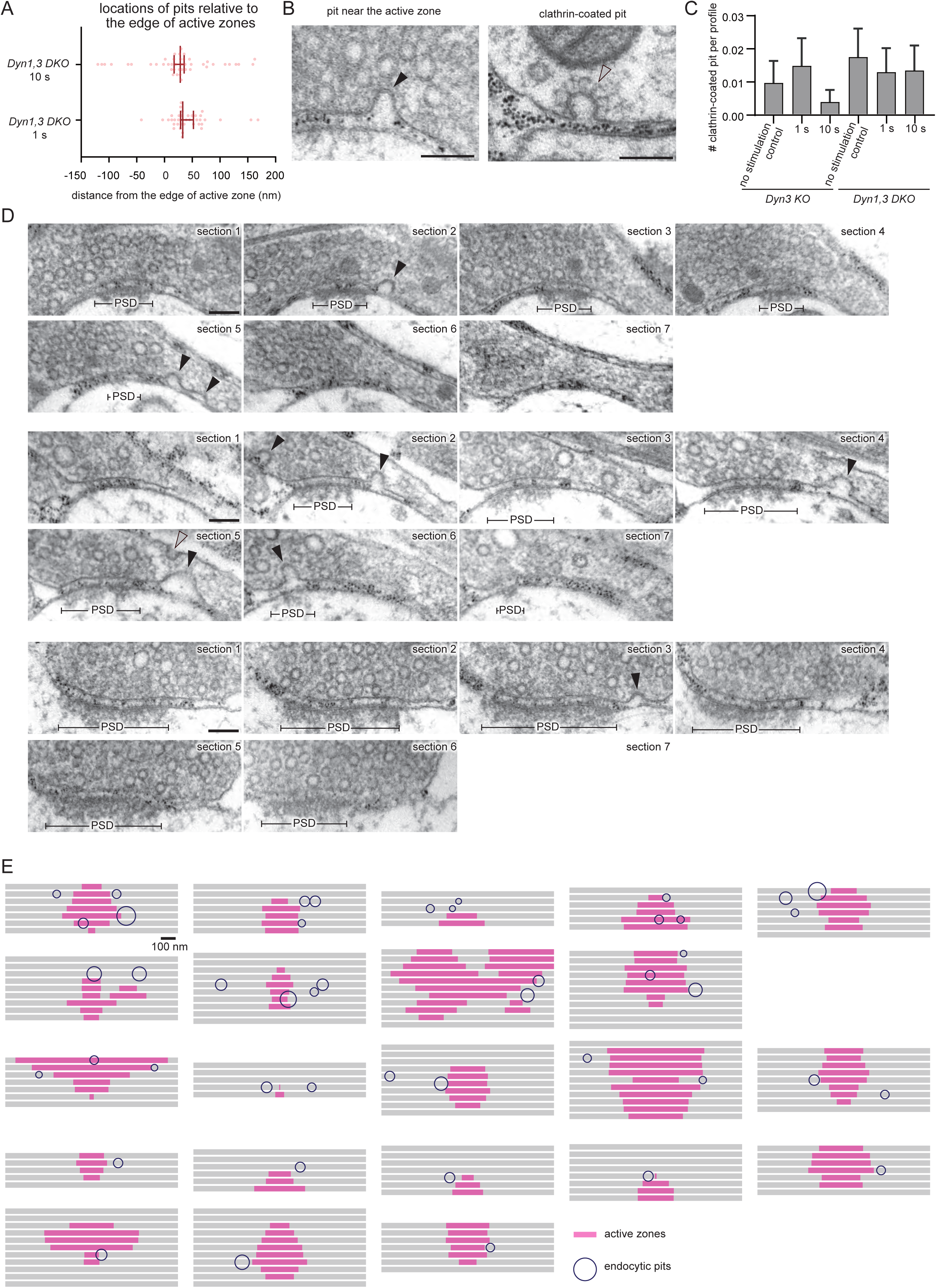
Endocytic pits in *Dyn1, 3* DKO neurons reconstructed from serial electron micrographs. (A) Locations of endocytic pits relative to the edge of an active zone. Each dot represents a pit. The negative and positive values indicate inside and out the active zone, respectively. The active zone is defined as the presynaptic membrane area juxtaposed to the postsynaptic density. The median and 95% confidence interval are shown. n = 57 pits for 1 s and 44 pits for 10 s. p = 0.2707. Mann-Whitney test was used. The same dataset as Figure 1. (B) Example transmission electron micrographs showing endocytic pits and clathrin-coated pit structures in *Dyn1, 3* DKO neurons. Scale bar: 100 nm. Black arrowheads, endocytic invaginations. Clear arrowheads, clathrin-coated pit. (C) Number of clathrin-coated pits in no stimulation control, at 1 s and at 10 s after stimulation. The numbers between *Dyn 3* KO and *Dyn1,3* DKO were not different, Ordinary one-way ANOVA with full pairwise comparisons by Tukey’s multiple comparisons test. The mean and SEM are shown. (D) Example transmission electron micrographs from serial sections of presynaptic terminals from Dyn1, 3 DKO neurons frozen 10 s after a single stimulus. Scale bar: 100 nm. PSD: post-synaptic density. Black arrowheads, endocytic invaginations. Translucent arrowheads, clathrin-coated pit. (E) Graphical depictions of serial-sectioned bouton containing endocytic pits around active zones, from the experiments described in (D).

**Figure S2.**
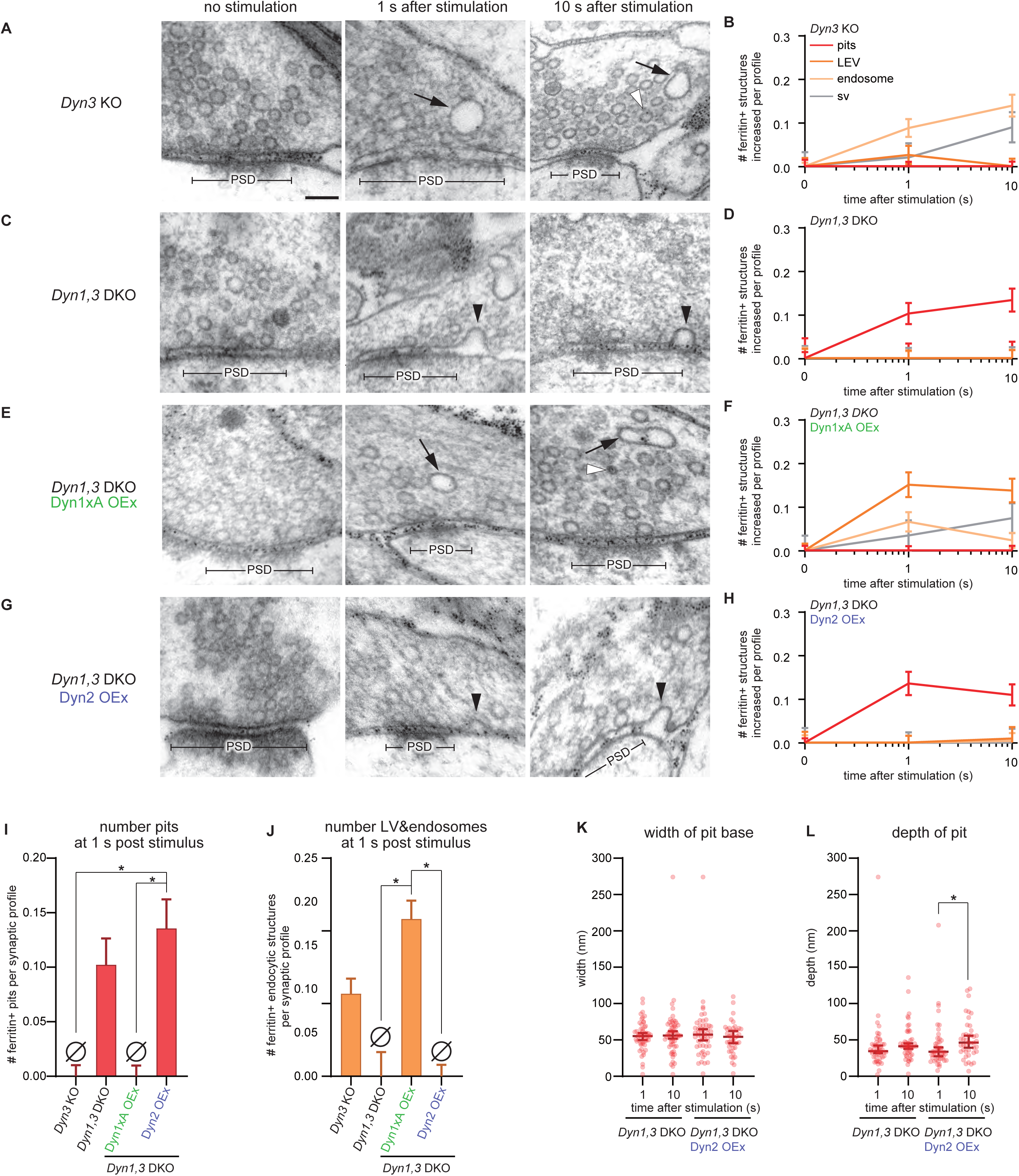
Dyn2 cannot rescue ultrafast endocytic defect of *Dyn1,3* DKO. (A, C, E and G) Example micrographs showing endocytic pits and ferritin-containing endocytic structures at the indicated time points in *Dyn3* KO neurons (A), *Dyn1, 3* DKO neurons (C), *Dyn1, 3* DKO neurons, overexpressing Dyn1xA (Dyn1xA OEx) (E) and *Dyn1, 3* DKO neurons, overexpressing Dyn2 (Dyn2 OEx) (G). Black arrowheads, endocytic pits; black arrows, ferritin-positive large endocytic vesicles (LEVs) or endosomes; white arrowheads, ferritin-positive synaptic vesicles. Note that Dyn2 OEx did no rescue the endocytic defect of *Dyn1, 3* DKO. Scale bar: 100 nm. PSD, post-synaptic density. (B, D, F and H) Plots showing the increase in the number of each endocytic structure per synaptic profile after a single stimulus in *Dyn3* KO neurons (B), *Dyn1, 3* DKO neurons (D), *Dyn1, 3* DKO, Dyn1xA OEx neurons (F) and *Dyn1, 3* DKO, Dyn2 OEx neurons (H). The mean and SEM are shown in each graph. (I) Number of endocytic pits at 1 s after stimulation. The numbers are re-plotted as a bar graph from the 1 s time point in (B, D, F and H) for easier comparison between groups. p = 0.0455 for *Dyn3* KO and 0.0455 for Dyn1xA OEx against Dyn2 OEx. Ordinary one-way ANOVA with full pairwise comparisons by Holm-Šídák’s multiple comparisons test. The mean and SEM are shown. (J) Number of ferritin-positive LEVs and endosomes at 1 s after stimulation. The numbers of LEVs and endosomes are summed from the data presented in (B, D, F and H), averaged, and re-plotted for easier comparison between groups. p = 0.0205 for *Dyn3* KO and 0.0205 for Dyn2 OEx against Dyn1xA OEx. Ordinary one-way ANOVA with full pairwise comparisons by Holm-Šídák’s multiple comparisons test. The mean and SEM are shown. (K and L) Plots showing the width (K) and depth (L) of endocytic pits at the 1s time point. The median and 95% confidence interval are shown in each graph. n = *Dyn1, 3* DKO, 51 pits for 1 s and 56 pits for 10 s; Dyn2 OEx, 52 pits for 1 s and 42 pits for 10 s. The depth: p = 0.0475 for Dyn2 OEx for 10 s against Dyn2 OEx for 1 s. Kruskal-Wallis Test with full comparisons by post hoc Dunn’s multiple comparisons tests. All data are from two independent experiments from N = 2 cultures prepared and frozen on different days. n = *Dyn* 3KO, 636; *Dyn1, 3* DKO, 631; *Dyn1, 3* DKO, Dyn1xA OEx, 617; and *Dyn1, 3* DKO, Dyn2, 600 synaptic profiles in (B, D, F, H, J, K, and L). *p < 0.05. See Quantification and Statistical Analysis for the n values and detailed numbers for each time point. Knock out neurons are from the littermates in all cases.

**Figure S3.**
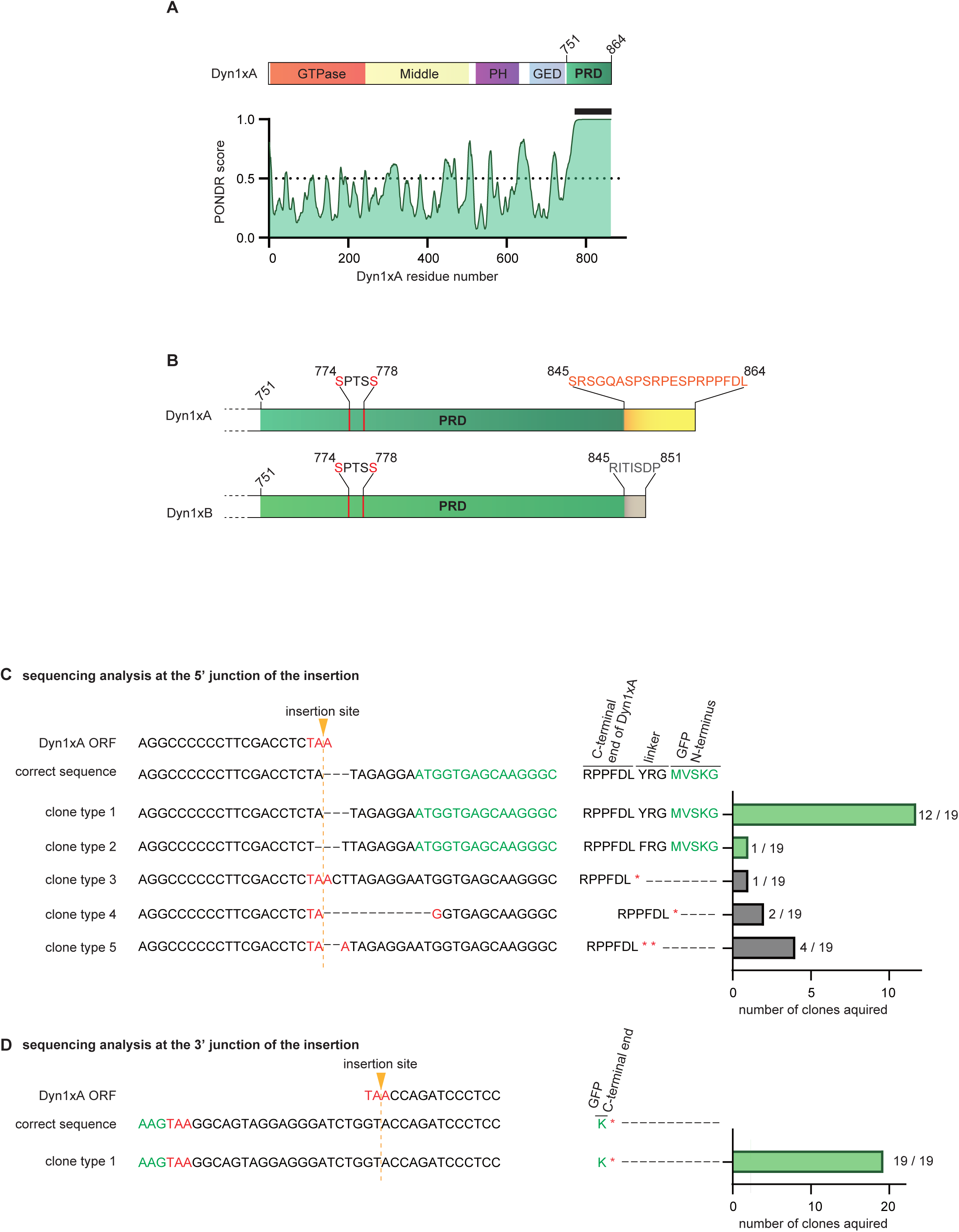
2D protein structures of Dyn1 and Sequencing analysis of CRISPR knock-in Dyn1xA. (A) A schematic showing the protein structural elements of Dyn1xA (top) and output of intrinsic disorder prediction program PONDR (http://www.pondr.com) for Dyn1xA (bottom). Dyn1xA contains the GTPase domain, the middle domain, the Pleckstrin homology (PH) domain, GTPase effector (GED) domain and Proline-rich domain (PRD). Only the PRD is the intrinsic disordered within Dyn1xA. An output of intrinsic disorder prediction program PONDR (http://www.pondr.com) for Dyn1xA (bottom). (B) A schematic showing the C-terminus of Dyn1xA and Dyn1xB. Both splice variants share S774/778 phosphorylation sites. Dyn1xA contains a 20 amino acid extension at the end of the C-terminus (yellow), whereas Dyn1xB has a calcineurin binding motif (PxIxIS) at the C-terminus (grey). P844 is shown in green. The calcineurin binding motif is underlined. (C) Sequencing analysis at the 5’ junction of the insertion site. The following sequences are shown: nucleotide sequences of the Dyn1xA open reading frame (ORF), an expected sequence after the CRISPR based knock-in (correct sequence), and 5 different clones acquired from cultured cortical neurons infected with virus containing the CRISPR knock-in construct. The number of each clone acquired is summarized on the left. Clone type 1; the expected sequence. Clone type 2; only 1 amino acid was substituted in the linker peptide between Dyn1xA and GFP. Clone types 3-5; codon shifts, causing a stop codon insertion after the Dyn1xA ORF. (D) Sequencing analysis at the 3’ junction of the insertion site. Only the correct clone was obtained from cultured cortical neurons infected with virus containing the CRISPR knock-in construct. The experiment was performed from the same neurons in (A).

**Figure S4.**
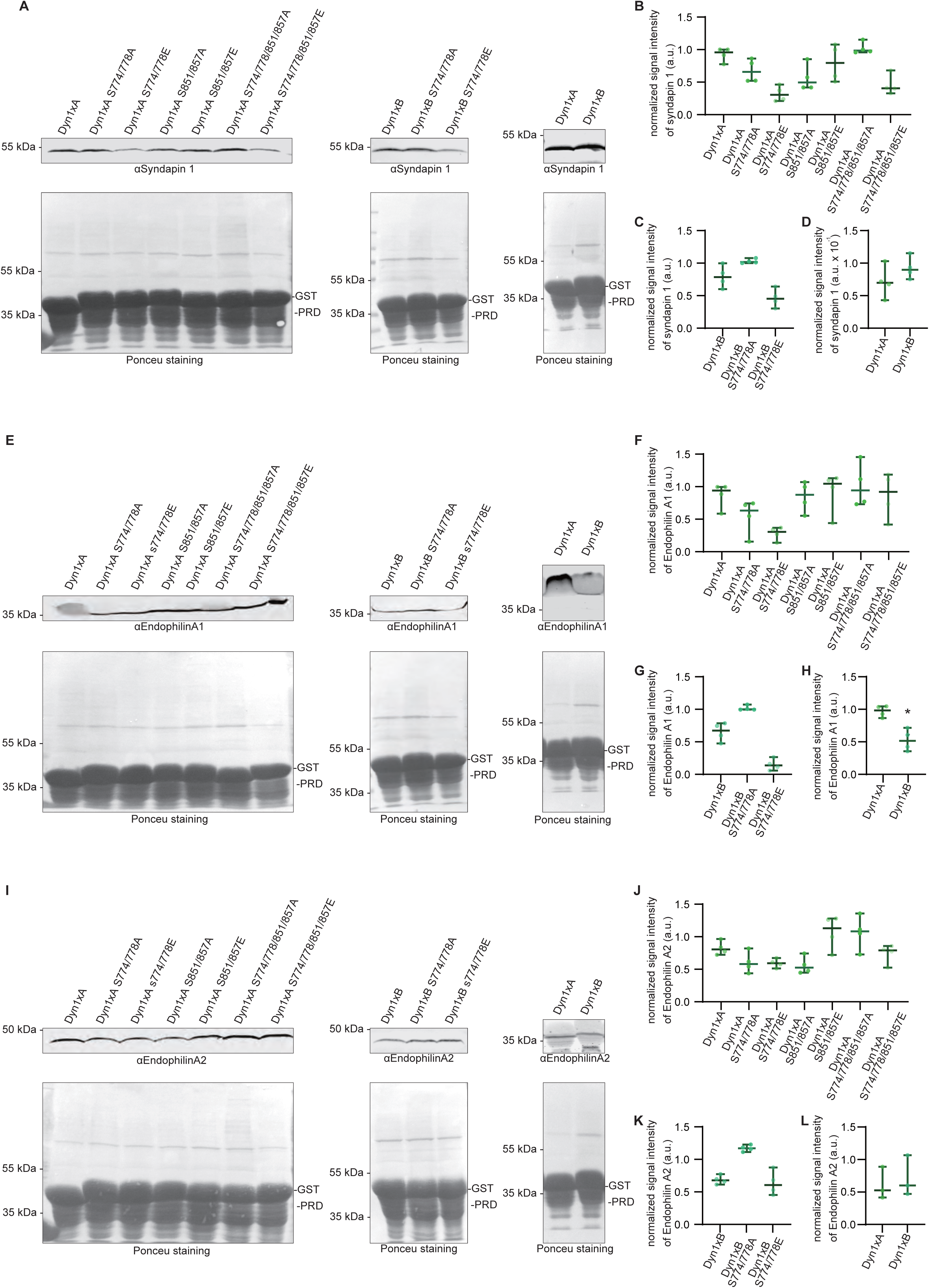
Pull-down assays of Dyn1xA binding partners. (A) The binding affinity of Dyn1xA-PRD (left) or Dyn1xB-PRD (middle) against Syndapin1. The binding affinity compared between Dyn1xA-PRD and Dyn1xB-PRD (right). Immunoblotting images showing a GST pull-down assay of the recombinant PRD domain of Dyn1xA, Dyn1xA S774/778A, Dyn1xA S774/778E, Dyn1xA S851/857A, Dyn1xA S851/857E, Dyn1xA S774/778/851/857A, Dyn1xA S774/778/851/857E, Dyn1xB, Dyn1xB S774/778A or Dyn1xB S774/778E, and anti-Syndapin 1 antibody reactions. The assay was performed using the lysate of synaptosome fractions isolated from adult mouse whole brains. Ponceu staining (bottom) showing the amount of the recombinant PRD of Dyn1xA or Dyn1xB constructs used for immunoblotting in each corresponding lane. (B) Normalized signal intensities of Syndapin 1 acquired from the immunoblotting using Dyn1xA PRD constructs shown in (A). Signal intensities were normalized to the amount of GST-PRDs based on the Ponceu staining. The median and 95% confidence interval are shown. (C) Normalized signal intensities of Syndapin 1 acquired from the immunoblotting using Dyn1xB-PRD constructs shown in (A). Signal intensities were normalized to the amount of GST-PRDs based on the Ponceu staining. The median and 95% confidence interval are shown. (D) Normalized signal intensities of Syndapin 1 between pull-down Dyn1xA-PRD and Dyn1xB-PRD acquired from the immunoblotting shown in (A). Signal intensities were normalized to the amount of GST-PRDs based on the Ponceu staining. N = 4 from different mouse brain. The median and 95% confidence interval are shown. p =0.4829 for Dyn1xB-PRD against Dyn1xA-PRD, unpaired t test. The median and 95% confidence interval are shown. (E) The binding affinity of Dyn1xA-PRD (left) or Dyn1xB-PRD (middle) against Endophilin A1. The binding affinity compared between Dyn1xA-PRD and Dyn1xB-PRD (right). Immunoblotting images showing a GST pull-down assay of the recombinant PRD domain of Dyn1xA, Dyn1xA S774/778A, Dyn1xA S774/778E, Dyn1xA S851/857A, Dyn1xA S851/857E, Dyn1xA S774/778/851/857A, Dyn1xA S774/778/851/857E, Dyn1xB, Dyn1xB S774/778A or Dyn1xB S774/778E, and anti-Endophilin A1 antibody reactions. The assay was performed using the lysate of synaptosome fractions isolated from adult mouse whole brains. Ponceu staining (bottom) showing the amount of the recombinant PRD of Dyn1xA or Dyn1xB constructs used for immunoblotting in each corresponding lane. (F) Normalized signal intensities of Endophilin A1 acquired from the immunoblotting using Dyn1xA PRD constructs shown in (E). Signal intensities were normalized to the amount of GST-PRDs based on the Ponceu staining. The median and 95% confidence interval are shown. (G) Normalized signal intensities of Endophilin A1 acquired from the immunoblotting using Dyn1xB-PRD constructs shown in (E). Signal intensities were normalized to the amount of GST-PRDs based on the Ponceu staining. The median and 95% confidence interval are shown. (H) Normalized signal intensities of Endophilin A1 between pull-down Dyn1xA-PRD and Dyn1xB-PRD acquired from the immunoblotting shown in (E). Signal intensities were normalized to the amount of GST-PRDs based on the Ponceu staining. N = 4 from different mouse brain. The median and 95% confidence interval are shown. p =0.0031 for Dyn1xB-PRD against Dyn1xA-PRD, unpaired t test. The median and 95% confidence interval are shown. (I) The binding affinity of Dyn1xA-PRD (left) or Dyn1xB-PRD (middle) against Endophilin A2. The binding affinity compared between Dyn1xA-PRD and Dyn1xB-PRD (right). Immunoblotting images showing a GST pull-down assay of the recombinant PRD domain of Dyn1xA, Dyn1xA S774/778A, Dyn1xA S774/778E, Dyn1xA S851/857A, Dyn1xA S851/857E, Dyn1xA S774/778/851/857A, Dyn1xA S774/778/851/857E, Dyn1xB, Dyn1xB S774/778A or Dyn1xB S774/778E, and anti-Endophilin A2 antibody reactions. The assay was performed using the lysate of synaptosome fractions isolated from adult mouse whole brains. Ponceu staining (bottom) showing the amount of the recombinant PRD of Dyn1xA or Dyn1xB constructs used for immunoblotting in each corresponding lane. (J) Normalized signal intensities of Endophilin A2 acquired from the immunoblotting using Dyn1xA PRD constructs shown in (I). Signal intensities were normalized to the amount of GST-PRDs based on the Ponceu staining. The median and 95% confidence interval are shown. (K) Normalized signal intensities of Endophilin A2 acquired from the immunoblotting using Dyn1xB-PRD constructs shown in (I). Signal intensities were normalized to the amount of GST-PRDs based on the Ponceu staining. The median and 95% confidence interval are shown. (L) Normalized signal intensities of Endophilin A2 between pull-down Dyn1xA-PRD and Dyn1xB-PRD acquired from the immunoblotting shown in (I). Signal intensities were normalized to the amount of GST-PRDs based on the Ponceu. N = 4 from different mouse brain. The median and 95% confidence interval are shown. p =0.6784for Dyn1xB-PRD against Dyn1xA-PRD, unpaired test. The median and 95% confidence interval are shown. N = 4 from different mouse brains in all cases. * p < 0.05. See Quantification and Statistical Analysis for the n values and detailed numbers.

**Figure S5.**
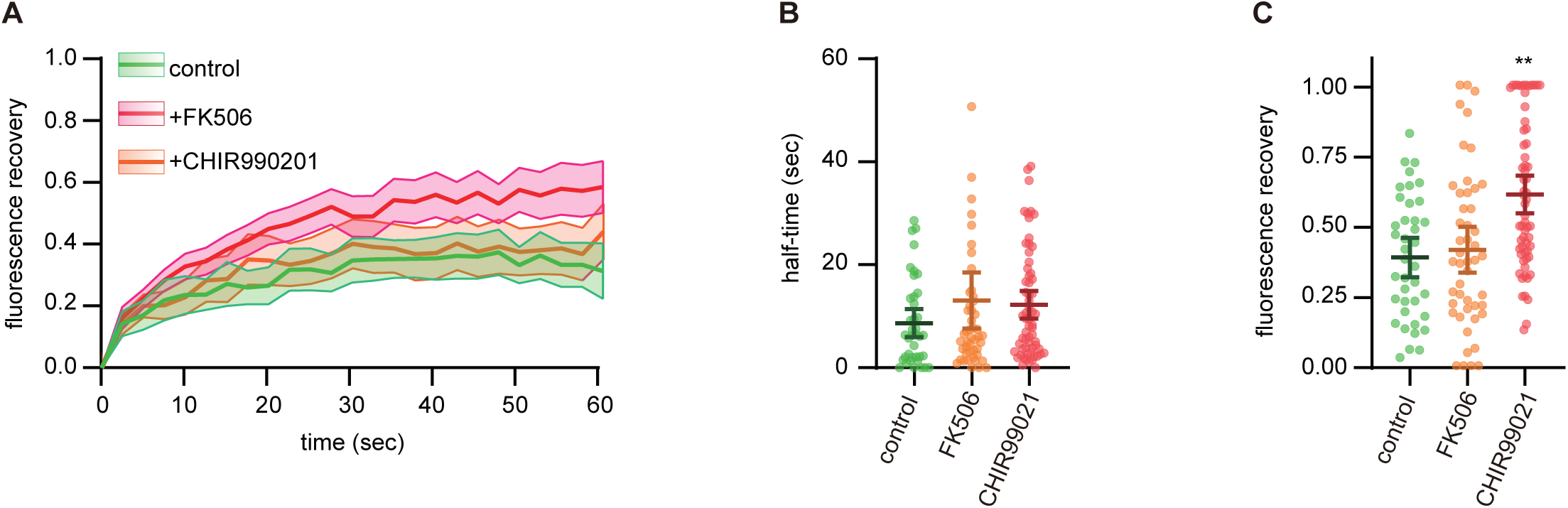
Dephosphorylation of PRD increases the mobile fraction of Dyn1xA. (A) Fluorescence recovery of Dyn1xA signals in DMSO (control), calcineurin inhibitor (FK506) or GSK3β inhibitor (CHIR99021) treated neurons. The entire puncta of Dyn1xA were photobleached in each case, and the recovery measured. The median and 95% confidence interval are shown. (B) Plots showing the half-time of Dyn1xA signals in the indicated conditions: control, FK506, CHIR99021. The median and 95% confidence interval are shown. The half-time is calculated at the recovery period of 60 s after photobleaching. The median and 95% confidence interval are shown. Each dot represents a punctum. The number was not different between the conditions. Kruskal–Wallis tests with comparisons against Dyn1xA by post hoc Dunn’s multiple comparisons tests. (C) Plots showing the fraction of fluorescence recovery 60 s after photobleaching of the entire Dyn1xA puncta in control, FK506, and CHIR99021-treated neurons. The median and 95% confidence interval are shown. Each dot represents a punctum. p = 0.0003 for CHIR99021 against control. Kruskal–Wallis tests with comparisons against Dyn1xA by post hoc Dunn’s multiple comparisons tests. n = Dyn1xA puncta, 71; FK506, 46; and CHIR99021, 64 in (A-C). At least 4 different neurons were examined from 2 independent cultures. The Dyn1xA puncta data are shared with Figure 3G-I. FK506 and CHIR99021 experiments were used same cultures as in Figure 3G-I.

**Figure S6.**
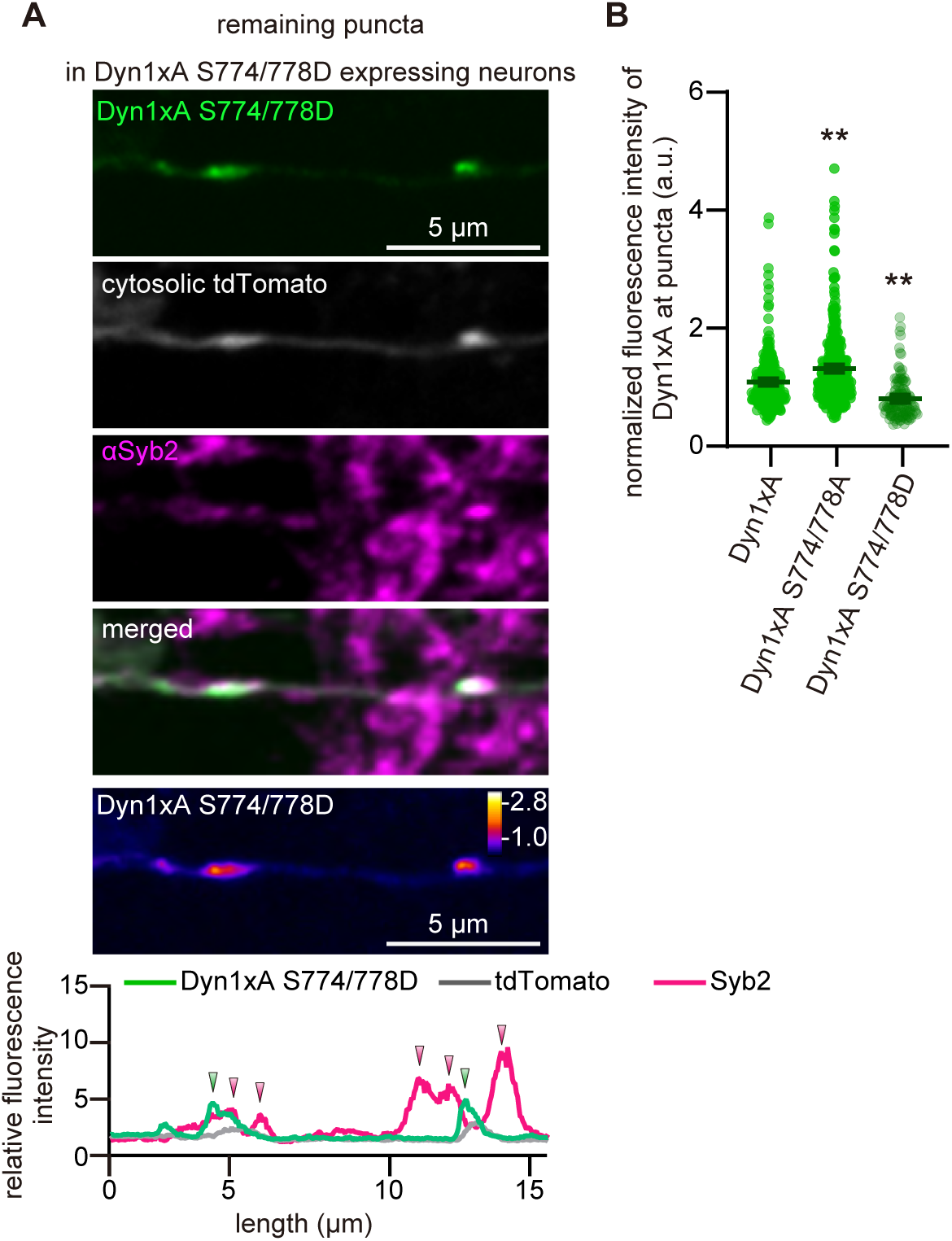
Quantification of remaining Dyn1xA S774/778D puncta. (A) Example confocal immunofluorescence micrographs showing the overexpression of Dyn1xA S774/778D along with exogenously expressed cytosolic tdTomato and immuno-stained *α*Syb2. False-colored images show the relative fluorescence intensity of Dyn1xA S774/778D. Line scan graphs represent the localization of Dyn1xA S774/778D relative to cytosolic tdTomato and *α*Syb2. (B) Normalized fluorescence intensities of Dyn1xA, Dyn1xA S774/778A or Dyn1xA S774/778D puncta. n = 287 puncta in Dyn1xA, 373 puncta in Dyn1xA S774/778A, and 123 puncta in Dyn1xA S774/778D from 2 separate cultures. The median and 95% confidence interval are shown. p < 0.0001 for Dyn1xA S774/778A and for Dyn1xA S774/778D against Dyn1xA. Kruskal–Wallis tests with comparisons against Dyn1xA by post hoc Dunn’s multiple comparisons tests. The same data as in Figure 4 are analyzed.

**Figure S7.**
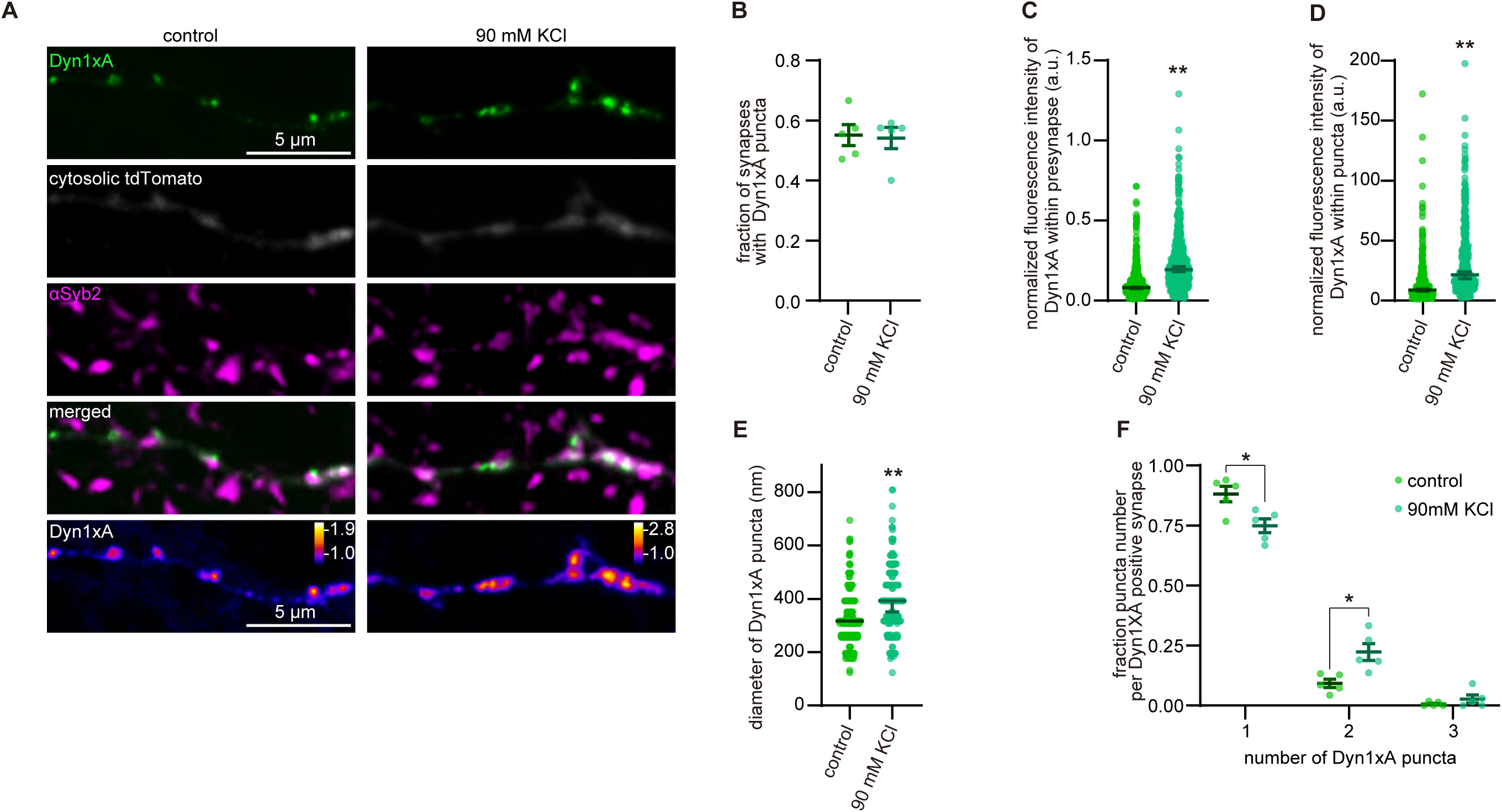
Synaptic activity controls the number of Dyn1xA puncta. (A) Example confocal fluorescence micrographs showing the localization of Dyn1xA, along with exogenously expressed cytosolic tdTomato and antibody staining of Synaptobrevin 2 (*α*Syb2) in fixed neurons 90 s after the control and the 90 mM KCl treatment. False-colored images of Dyn1xA (bottom panels) show the relative fluorescence intensity of these molecules. (B) The fraction of presynapses that contain Dyn1xA puncta 90 s after sham (2.5 mM KCl) or the 90 mM KCl treatment. n = 5 neurons in each condition. Each dot represents a fraction calculated from each neuron. (C) The normalized fluorescence intensities of Dyn1xA within presynapses 90 s after sham or the 90 mM KCl treatment. n = 674 in the control and 586 in the 90 mM KCl treatment. The median and 95% confidence interval are shown. p = 0.732, Mann-Whitney test. Each dot on the plot represents a punctum. (D) The normalized fluorescence intensities of Dyn1xA within puncta 90 s after sham or the 90 mM KCl treatment. n = 414 in the control and 402 in the 90 mM KCl treatment. The median and 95% confidence interval are shown. p < 0.0001, Mann-Whitney test. Each dot on the plot represents a punctum. (D) Diameters of Dyn1xA puncta 90 s after sham or the 90 mM KCl treatment. n = 414 in the control and 402 in the 90 mM KCl treatment. The median and 95% confidence interval are shown. p < 0.0001, Mann-Whitney test. Each dot on the plot represents a punctum. (F) Relative frequency distributions of the number of puncta within presynaptic boutons among those that contain at least one punctum. The fraction is calculated from each neuron. The mean and SEM are shown. n = 5 neurons in each condition. p = 0.0066 for 1 punctum and 0.0064 for 2 puncta in the KCl treatment against the control. Two-way ANOVA with Šídák’s multiple comparisons test. N = 2 independent cultures. *p < 0.05, **p < 0.0001. See Quantification and Statistical Analysis for the n values and detailed numbers.

**Figure S8.**
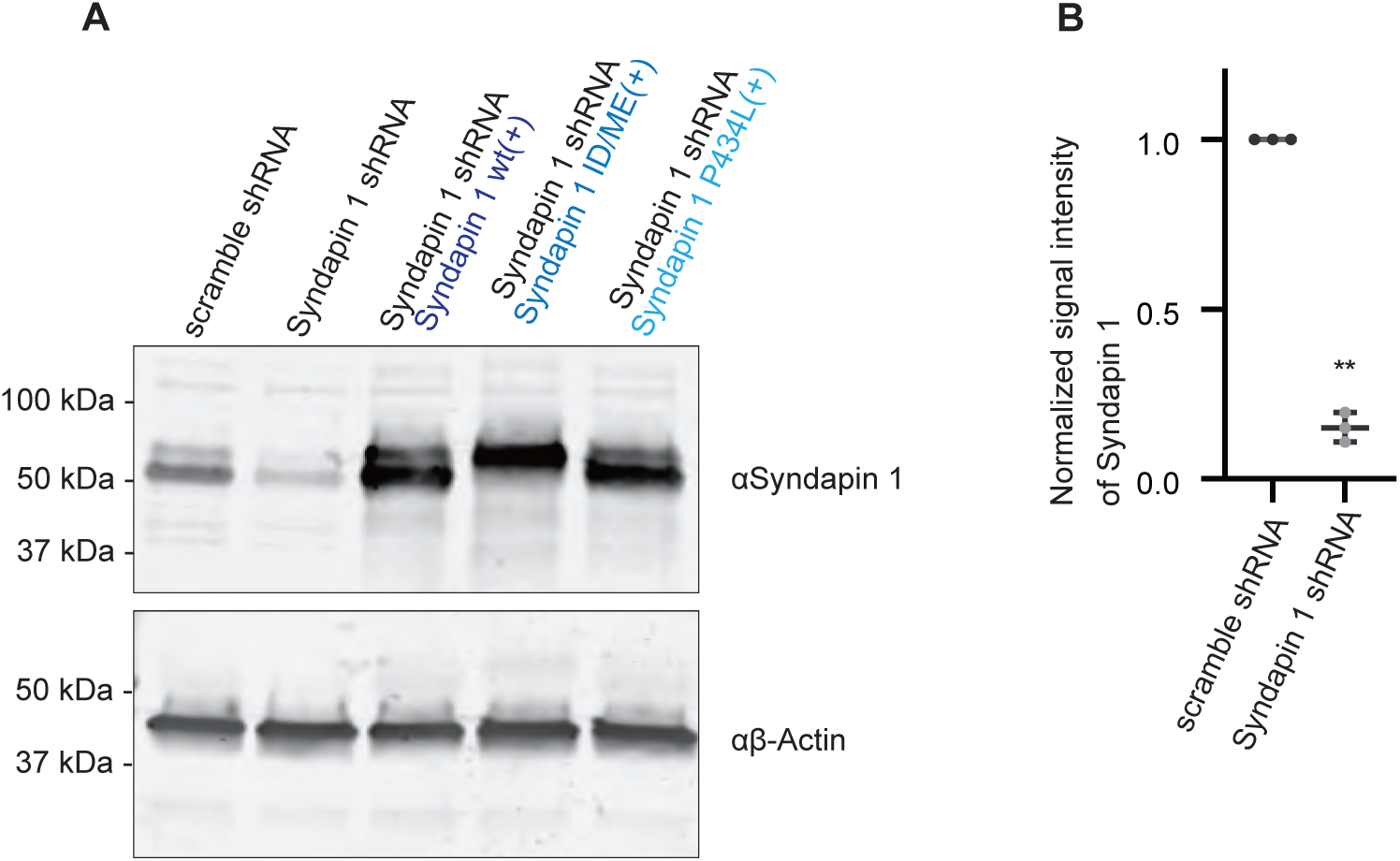
Evaluation of Syndapin 1 shRNA efficiency. (A) Immunoblotting images showing anti-Syndapin 1 or anti-β-Actin antibodies reactions against the lysates of cultured hippocampal neurons infected with lentivirus expressing scramble shRNA, Syndapin 1 shRNA, Syndapin 1 shRNA and Syndapin 1 wild type (wt) (+), Syndapin 1shRNA + Syndapin 1 I122D/M123E (ID/ME) (+) or Syndapin 1shRNA + Syndapin 1 P434L (+). (B) Normalized signal intensities of Syndapin 1 in immunoblotting assays using neurons expressing scramble shRNA or Syndapin shRNA. **p < 0.0001, unpaired t test. The mean and SEM are shown. n = 3 independent cultures.

